# Association of milk microbiome in bovine clinical mastitis and their functional implications in cows in Bangladesh

**DOI:** 10.1101/591982

**Authors:** M. Nazmul Hoque, Arif Istiaq, Rebecca A. Clement, Munawar Sultana, Keith A. Crandall, Amam Zonaed Siddiki, M. Anwar Hossain

## Abstract

Milk microbiomes impose a significant influence on the pathophysiology of bovine mastitis. To assess the association, we compared the microbiome of clinical mastitis (CM) and healthy (H) milk samples through whole metagenomic deep sequencing. A total of 483.38 million reads generated from both metagenomes were analyzed through PathoScope and MG-RAST, and mapped to 380 bacterial, 56 archaeal, and 39 viral genomes. We observed distinct shifts and differences in abundance between the microbiome of CM and H milk in phyla Proteobacteria, Bacteroidetes, Firmicutes and Actinobacteria with an inclusion of 68.04% unreported and/or opportunistic species in CM milk. Additionally, 14 archaeal and 14 viral genera were found to be solely associated with CM. The functional metagenomics identified several pathways related to bacterial proliferation and colonization such as metabolism, chemotaxis and invasion, immune-diseases, oxidative stress, regulation and cell signaling, phage and prophases, antibiotic and heavy metal resistance to be associated with CM. Therefore, the present study provides conclusive data on milk microbiome diversity associated with bovine CM and its role in udder health.

## Introduction

Mastitis is one of the most prevalent diseases in the dairy industry with the highest clinical and economic significance worldwide^1^. The condition usually happens when pathogenic microbes enter the germ-free environment of the mammary gland, mostly by the disruption of the physical barriers of the mammary quarters, requiring prompt and appropriate host defenses to prevent colonization and subsequent disease pathology^2^. Diverse groups of microbes are known to colonize the mammary quarters of cows, and have evolved novel mechanisms that facilitate their proliferation, leading to clinical mastitis (CM). Despite knowledge of a few of these invading microbial groups, the etiology of bovine mastitis is continuously changing, with new microbial species identified as causing disease frequently. Additionally, although bacteria are the main cause of mastitis^3^, other microbes like archaea, viruses and fungi might be associated with the disease process^4^ and should therefore be investigated as well. During the progression of the mastitis, dysbiosis of the milk microbiome can occur with the increase of opportunistic pathogenic bacteria and reduction of healthy commensal bacteria^5^. Until recently, investigations of microbiome associated with bovine mastitis have been mostly restricted to individual pathogen isolation and characterization.

The disease is caused by epidemiologically diverse groups of microorganisms and categorized into contagious and environmental mastitis^6^. The udder of the dairy cows is the primary reservoir of contagious pathogens including *Staphylococcus aureus, Streptococcus agalactiae, Streptococcus dysgalactiae, Mycoplasma* spp. and *Corynebacterium bovis*^1, 6^. The involvement of the bovine mammary gland microbiota in the host-pathogen interaction has little investigated except during the infectious episode^7^. Environmental pathogens such as *Escherichia coli, Klebsiella pneumoniae, Klebsiella oxytoca, Enterobacter aerogenes, Streptococcus dysgalactiae* and *Streptococcus uberis*^1, 6^ can also be implicated in disease. Rapid advances in high-throughput NGS technology and bioinformatics tools^8^ during the last decade have initiated a transition from clinical microbiology to genomic characterization of the microbiome associated with infection, including mastitis in lactating women^5^ and animals^9^. Shotgun whole metagenome sequencing (WMS) produces a metagenome reflecting the total microbial makeup of a sample (pathogenic, environmental, bacterial, fungal, viral) and has been used successfully to gain insights into the phylogenetic composition and species diversity of a variety of microbiomes^10^, including profiling of their functional attributes^11^. Thus, data can be generated regarding the identity and abundance of genes related to microbial metabolism, virulence and antibiotic resistance enabling identification of unknown etiological agents that play a role in mammary gland pathogenesis.

Overexpression of putative genes encoding immune suppression^12^, systemic oxidative stress^3^, and inflammatory processes^13^ are the crucial factors affecting the progression of CM. Indiscriminate and overuse of antibiotics to treat mastitis is main cause of multidrug resistant bacteria^14^. Therefore, summarizing the variation in biota and protein functional diversity in clinical and healthy milk microbiomes using cutting-edge genomic technologies like WMS^15^ and associated bioinformatic tools is essential to understanding the pathophysiological conditions of bovine CM. Here we report the first study of its kind where high-throughput sequencing data (on an average 23.01 million reads per sample) were generated to investigate the microbiome of bovine CM and H milk^16^. The results revealed that cows suffering from CM milk had a distinct microbial community with reduced diversity, higher relative abundance of opportunistic pathogens, and altered protein functions compared to their healthy counterparts.

## Results

### Structure and composition of the bovine milk microbiome

Compared to healthy (H) microbiomes, clinical mastitis (CM) milk microbiomes showed significantly reduced Shannon-estimated microbial richness (H; *p*=0.005, CM; *p*=0.007, U test). Species richness in both metagenomes also differed significantly between two bioinformatics tools (PS; *p*=0.039, MR; *p*=0.001, U test) (Supplementary Fig. 1). Beta diversity (PCoA) revealed significant microbial disparity (*p*=0.001) between CM and H sample groups (Supplementary Fig. 2). At phylum level, NMDS showed distinct diversity differences between the sample categories (Supplementary Fig. 3).

At the domain level, bacteria were the most abundant community, with an average abundance of 98.00%, followed by eukaryotes (1.80%), archaea (0.02%), viruses (0.04%), and unassigned sequences (0.002%) (Supplementary Data 1). Though the relative abundance of microbes was higher in CM compared to H milk, the abundance fluctuated more (CV=886.90 vs 511.80; PS, CV= 1521.41 vs 1221.92; MR). The unique and shared distribution of microbial taxa found in CM and H samples by two analytic tools is represented in Venn diagrams (Fig.1). A total of 363 bacterial species in CM, and 146 species in H metagenomes were detected in PS analysis, of which 116 (29.51%) species shared in the both conditions (Fig. 1a). However, through MR pipeline, 356 and 251 bacterial genera were detected in CM and H samples respectively, whereas 227 (63.8%) genera were common in both metagenomes (Fig. 1b). By comparing the detected bacterial genera between two analytic tools, 98 unique genera were identified, of them 62.24% genera were solely associated with the onset of bovine CM (Fig. 1c, Supplementary Data 2). In addition, MR detected 54 and 42 archaeal, and 35 and 25 viral genera, respectively in CM and H samples, and among them 25.00% and 35.00% archaeal and viral genera respectively had sole association with CM (Fig. 1d, e). Unlike MR, PS detected only one archaeal genera (*Methanobrevibacter*) in CM and none in H samples.

**Fig. 1.**
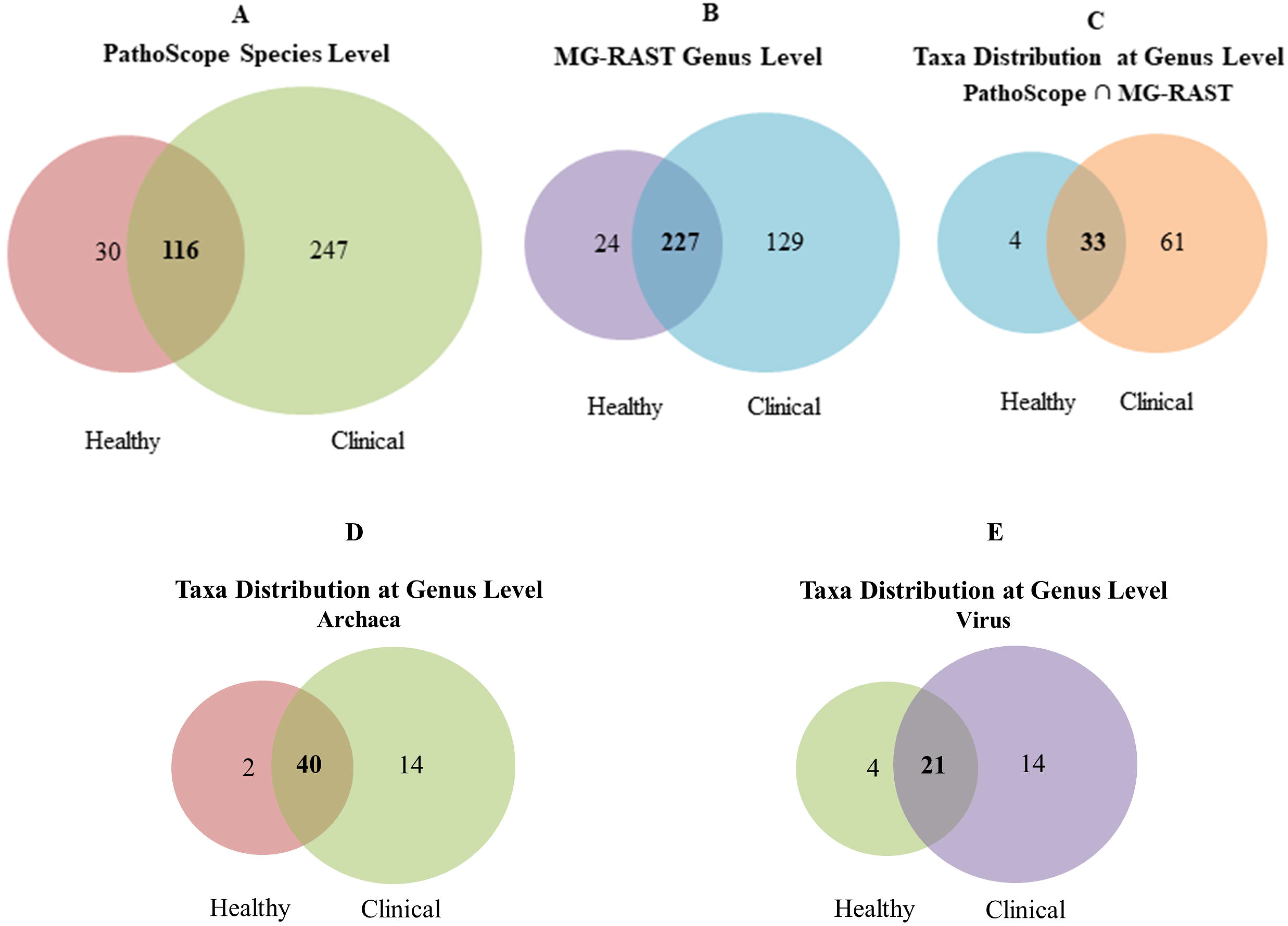
Venn diagrams representing the core unique and shared microbiomes of bovine clinical mastitis (CM) and healthy (H) milk. **a** Top Left: Venn diagram comparison of bacteria at strain level by PathoScope (PS), **b** Top Middle: Venn diagram showing unique and shared bacterial genera by MG-RAST (MR), **c** Top Right: Shared and unique bacterial genera distribution between PS and MR, **d** Bottom Left: & **e** Bottom Right: Venn diagrams representing unique and shared Archaeal and viral genera respectively found in bovine milk as analysed with MR pipeline. Microbiotas sharing between the conditions are indicated by bold colored.

### CM-associated bacteria changes at the genus level

The current microbiome study demonstrated notable differences among the microbial community in CM and H milk in both bioinformatics tools. Proteobacteria, Bacteroidetes, Firmicutes and Actinobacteria (contributing to 96.51% of the total sequences, U test, *p*=0.001) were the four most abundant phyla in PS and MR analyses. The relative abundance of the top 40 bacterial genera was compared between CM and H cohort through PS (Fig. 2) analyses. Among the predominating phyla, Proteobacteria was among the most diverse, and included a wide variety of genera including *Acinetobacter*, *Pseudomonas*, *Escherichia*, *Vibrio*, *Erwinia*, *Pantoea*. The phylum Firmicutes was dominated by *Streptococcus*, *Enterococcus*, *Staphylococcus*, and *Bacillus* genera while *Chryseobacterium*, *Porphyromonas* and *Prevotella* genera were predominating in Bacteroidetes phylum, and *Corynebacterium* was the most abundant genus in phylum Actinobacteria. Among the detected genera either of the tool, *Acinetobacter* (60.14%), *Campylobacter* (10.93%), *Pantoea* (0.66%), *Klebsiella* (0.63%), *Kluyvera* (0.42%), *Salmonella* (0.31%), *Enterobacter* (0.30%), *Shewanella* (0.30%), *Escherichia* (0.28%), *Citrobacter* (0.15%) and *Bacillus* (0.10%) had higher mean relative abundance in CM samples in both analytic tools, while rest of the genera had relatively lower mean abundance (<0.10%). In contrast, the H milk metagenomes also had higher mean relative abundance of genus *Acinetobacter* (52.90%) in PS and MR followed by *Pseudomonas* (22.81%), *Micromonospora* (10.57%), *Eubacterium* (5.37%), *Catenibacterium* (2.12%) and *Ralstonia* (0.12%)genera, and rest of the genera had much lower abundance (<0.10%). In general, MR detected higher number of microbial genera than PS (Supplementary Tables 2&3), however results from the both tools were concordant, with 98.00% of the total microbial abundance composed of shared genera (Supplementary Table 4, Supplementary Data 2).

**Fig. 2.**
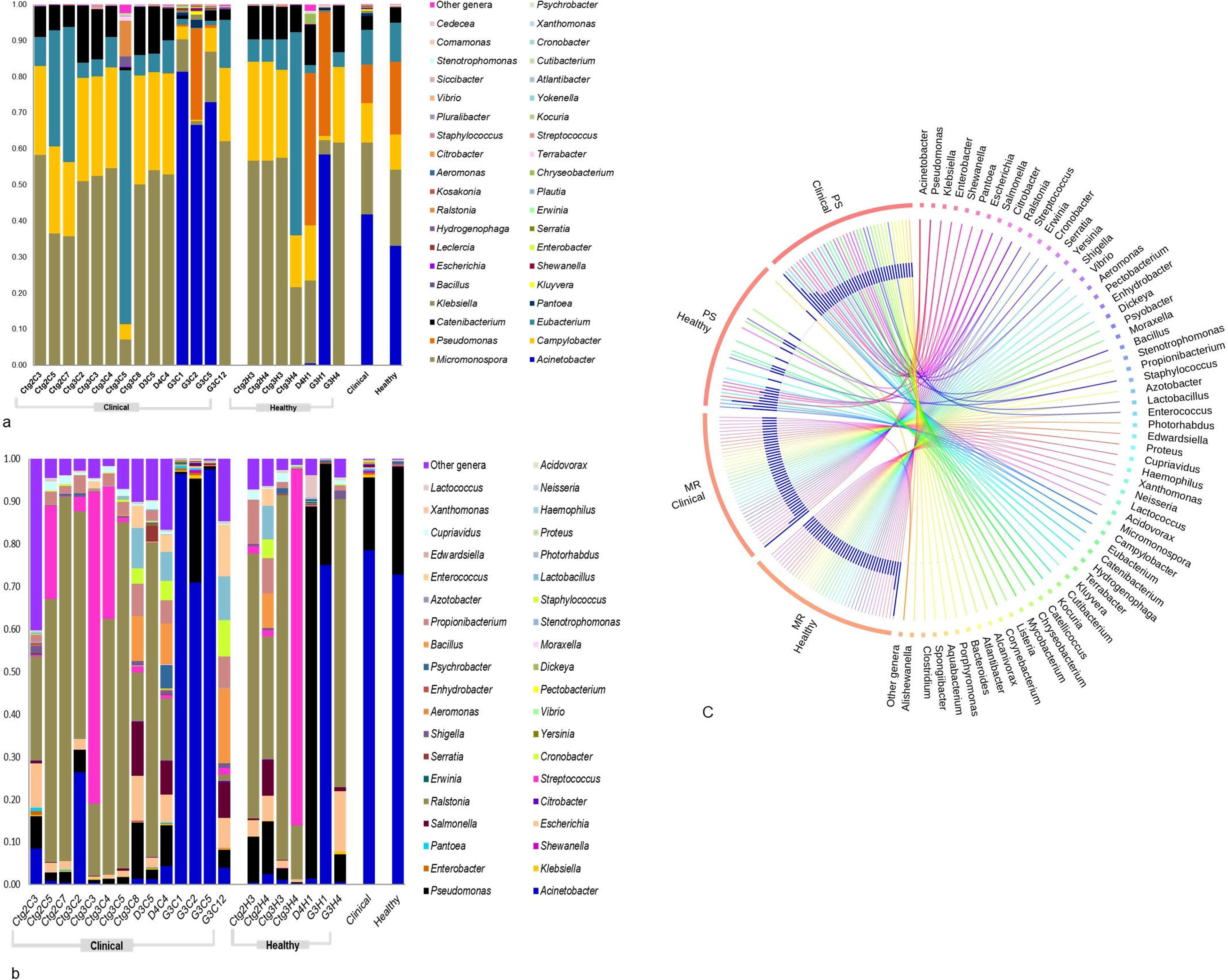
Taxonomic profile of 40 most abundant bacterial genera in bovine clinical mastitis (CM) and healthy (H) milk samples. **a** Top Left: abundance through PathoScope (PS) and **b** Bottom Left: through MG-RAST (MR) analyses. The 39 most abundant genera are sorted by descending order of the relative abundance in 21samples, with the remaining genera grouped into the ‘Other genera’. Each stacked bar plot represents the abundance of bacteria in each sample of the corresponding category, where the last two bar plots depict overall relative abundance of bacterial genera between CM and H samples, respectively. **c** Middle Right: The circular plot illustrates the relative abundance of top 40 bacterial genera in CM and H milk samples analysed through PS and MR. Taxa in both metagenomes are represented by different colored ribbons both tools. The relative abundancies are illustrated by the sizes of each color segment in the outer circle and the inner blue colored bars. Part of the microbiome is shared by both sample categories (CM-H milk), and part is analytic tool specific (PS-MR). Notable differences between the bacterial populations are those where the taxon is abundant in CM samples and effectively undetected in the H milk. Sample names: suffix ends with C refers to clinical (CM) and that ends with H refers to healthy (H) milk samples.

### CM-associated bacteria changes at the strain level

We further investigated whether strain level relative abundances of the bacteria differed between CM and H samples (Fig. 3, 4). The CM milk metagenome had significantly (*p*=0.001) higher number of bacterial species than the H milk, and among the detected species 62.85% had unique association with bovine CM, and 7.63% were solely found in H milk (Fig. 1a). The presence of few predominating bacterial species in both categories of samples suggests that the crucial differences might be occurring at the strain level, and most of the species identified in each sample were represented by a single strain. The CM milk metagenome was dominated by 26 strains (7.16%) of *Acinetobacter* species while *Pseudomonas*, *Streptococcus*, *Corynebacterium*, *Staphylococcus*, *Enterococcus*, *Bacillus*, and *Escherichia* species were represented respectively by 22, 16, 12, 11, 8, 7 and 6 different strains. However, in both metagenomes, *Acinetobacter johnsonii* XBB1 remained as the most abundant strain with a relative abundance of 39.03% and 31.23% respectively in CM and H samples. The other predominant strains in CM metagenome were *Campylobacter mucosalis*, *Bacillus mycoides*, *Klebsiella pneumoniae* subsp. pneumoniae HS11286, *Leclercia adecarboxylata*, *Escherichia coli* str. K-12 substr. MG1655, *Escherichia coli* O157:H7 str. Sakai, *Escherichia coli* UMN026, *Escherichia coli* IAI39, *Staphylococ cusaureus* subsp. aureus NCTC 8325, *Staphylococcus xylosus*, *Bacillus subtilis* subsp. subtilis str. 168, *Mycobacterium* sp. Root 265, *Macrococcus caseolyticus*. Importantly, this study demonstrated that 68.04% of the detected bacterial strains were exclusively found in CM milk metagenome, and among them *Pantoea dispersa* EGD-AAK13, *Klebsiella oxytoca, Kluyvera intermedia, Shewanella oneidensis* MR-1, *Kluyvera ascorbata* ATCC 33433, *Klebsiella aerogenes* KCTC 2190, *Kluyvera cryocrescens* NBRC 102467, *Acinetobacter pittii* PHEA-2, *Pseudomonas mendocina* ymp, *Acinetobacter gyllenbergii* NIPH 230 were the most predominant strains. Furthermore, most of these strains were previously unreported and possibly played an opportunistic role in the mammary gland pathogenesis (Supplementary Data 2, Supplementary Table 5).

**Fig. 3.**
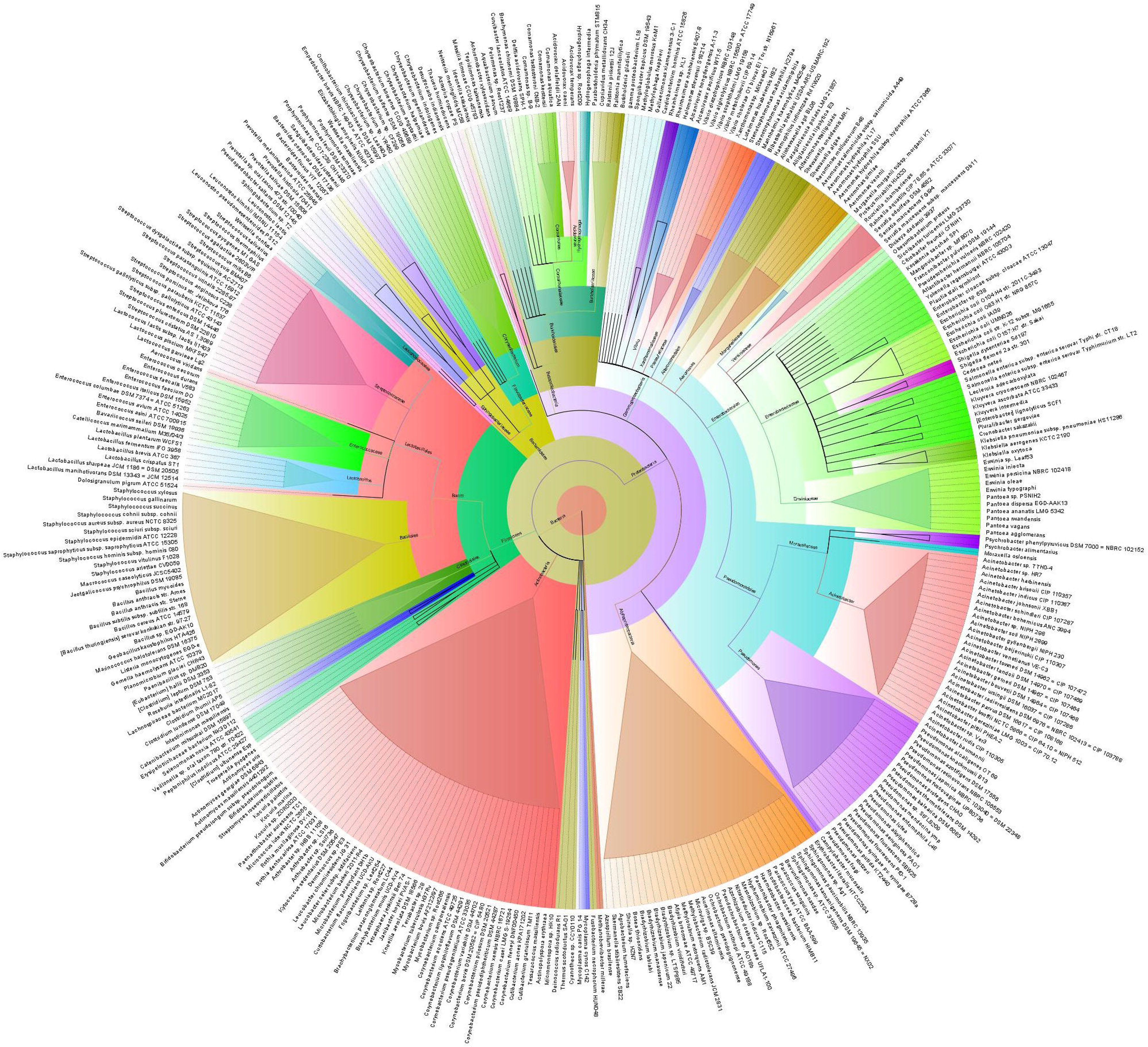
Taxonomic representation of unique microbiota associated to the bovine clinical mastitis (CM) milk at strain level. Sequences are assigned to different taxonomic index in PathoScope analysis using minimum identity of 95% and minimum alignment length 20 as cutoff parameters, and the circular phylogenetic tree is constructed based on the neighbor-joining algorithm using FigTree. The round tree illustrates 363 unique strains of bacteria in CM milk metagenomes. The inner circle represents the root of the microbiome defined as bacteria present in all samples. The outer circles represent different strains of bacteria is defined as species (with different strains) present in >50% of samples of the corresponding groups. For the outer circles, the width of a segment is proportional to the observed incidence for that species. Different colors are assigned according to the taxonomic ranks of the bacteria. The species and/or strains in the phylogenetic tree are also available in supplementary Data 2.

**Fig. 4.**
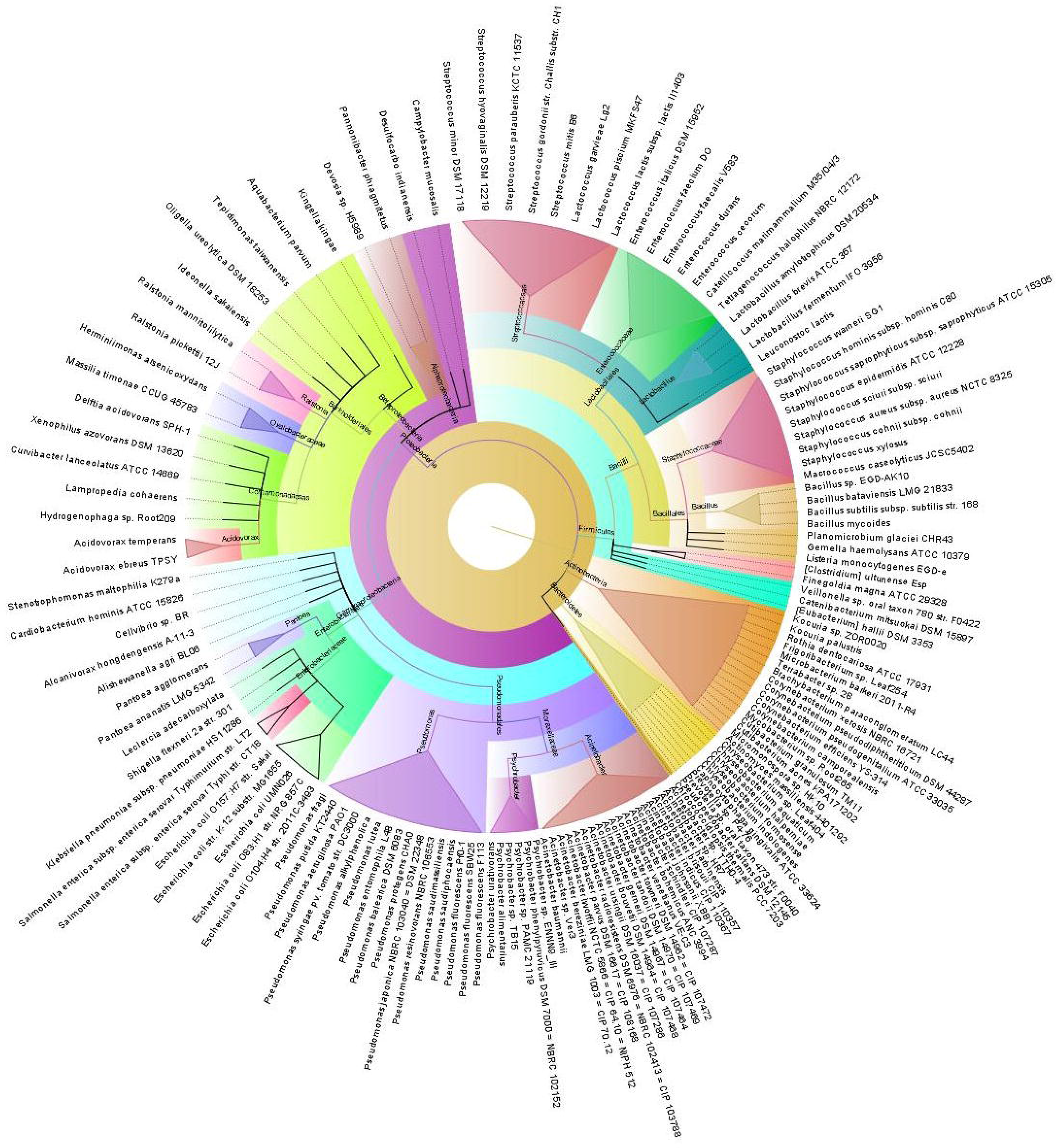
Taxonomic representation of unique microbiota associated to the bovine clinical mastitis (CM) and health (H) milk at strain level. Sequences are assigned to different taxonomic index in PathoScope analysis using minimum identity of 95% and minimum alignment length 20 as cutoff parameters, and the circular phylogenetic tree is constructed based on the neighbor-joining algorithm using FigTree. The round tree illustrates 146 unique strains of bacteria in H milk metagenomes. The inner circle represents the root of the microbiome defined as bacteria present in all samples. The outer circles represent different strains of bacteria is defined as species (with different strains) present in >50% of samples of the corresponding groups. For the outer circles, the width of a segment is proportional to the observed incidence for that species. Different colors are assigned according to the taxonomic ranks of the bacteria. The species and/or strains in the phylogenetic tree are also available in supplementary Data 2.

### CM-associated changes of archaea and viruses at the genus level

Archaea and viruses were detected in the samples of the both metagenomic groups; CM and H milk. The CM metagenome was dominated by *Methanosarcina* (41.94%), *Methanococcoides* (19.58%), *Methanococcus* (12.30%), *Methanocaldococcus* (2.59%), *Methanobrevibacter* (1.85%), *Thermococcus* (1.79%), and *Methanosphaera* (1.53%) archaeal genera with a lower relative abundance (<0.05%) of the rest of the genera (Fig. 5 a, Supplementary Data 2). Interestingly, none of the archaeal genus was detected in one CM sample (Ctg3C2). In contrast, *Methanoplanus* (14.69%), *Methanoculleus* (12.85%), *Euryarchaeota* (4.67%), and *Haloarcula* (1.50%) were the most abundant archaeal genera in H samples. The viral fraction of the current bovine milk microbiome was largely dominated by the members of the Caudovirales order, represented by the Podoviridae, Siphoviridae, and Myoviridae families. The predominating viral genera found in CM were *Epsilon15-like viruses* (15.78%), *P2-like viruses* (10.12%), *Myovirus* (8.18%), *Lambda-like viruses* (8.06%), *Bpp-1-like viruses* (7.12%), *phiKZ-like viruses* (4.35%), *Betaretrovirus* (2.01%), *P1-like viruses* (1.79%) and *T4-like viruses* (1.79%). The H milk however had relatively higher abundance of *Siphovirus* (55.85%), *Podovirus* (12.49%), *T1-like viruses* (3.44%) and *P22-like viruses* (1.71%) (Fig. 5 b, Supplementary Data 2).

**Fig. 5.**
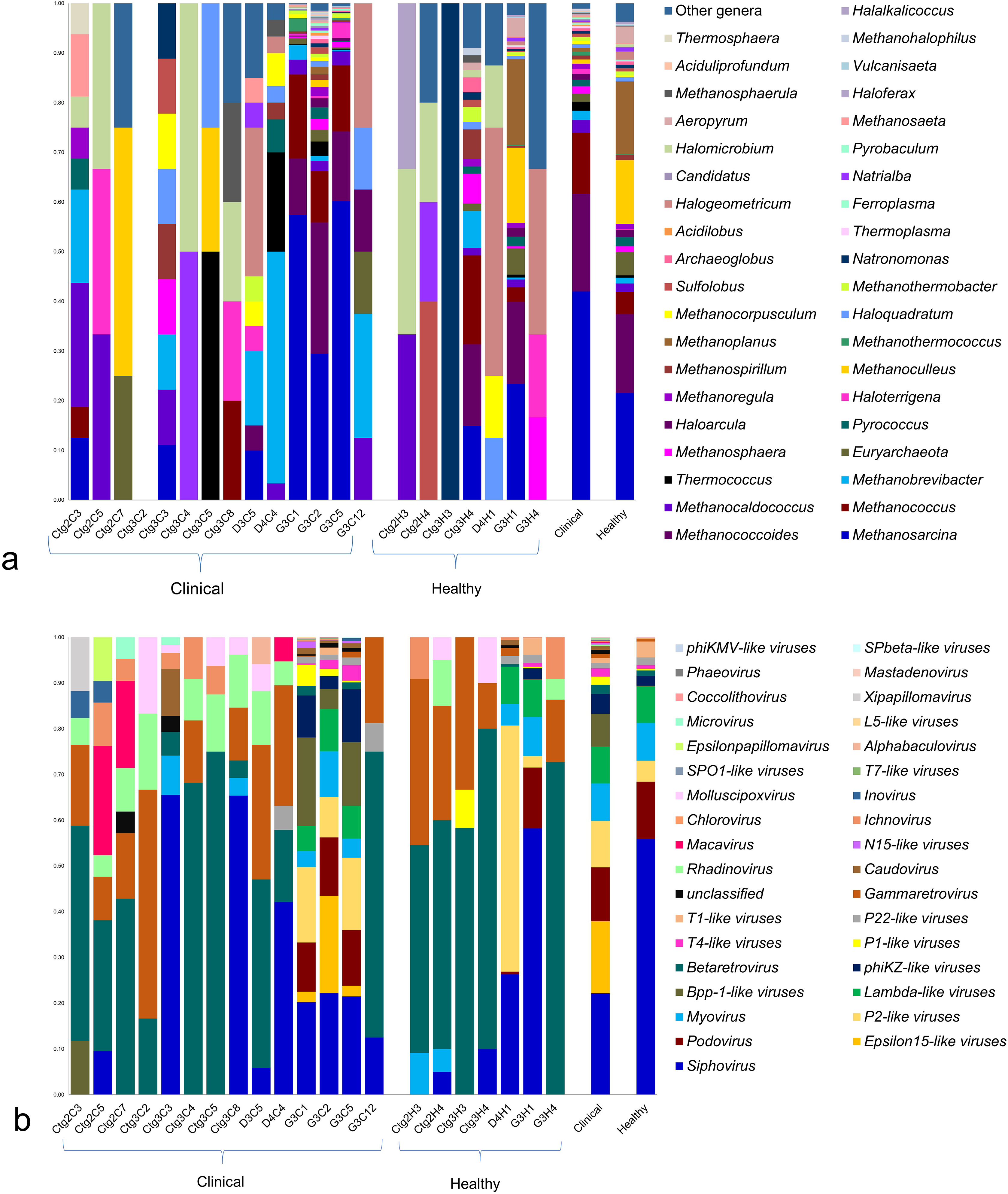
Taxonomic abundance of top 40 archaeal and viral genera from the reads count of MR output. **a** Top: Archaeal genera are found in 20 samples, and absent in one clinical sample (Ctg3C2). The 39 most abundant archaeal genera are sorted by descending order of the relative abundance, with the remaining genera keeping into the ‘Other genera’. **b** Bottom: Taxonomic distribution of 35 viral genera detected in all of the 21 samples of clinical (CM) and healthy (H) milk metagenomes. The most abundant viral genera are sorted by descending order of the relative abundance. Each stacked bar plot represents the abundance of archaea and viruses in each sample of the corresponding category, where the last two bar plots depict overall relative abundance of archaeal and viral genera in both metagenome groups. Notable differences between the archaeal and viral populations are those where the taxon is abundant in clinical samples and effectively undetected in the healthy milk. Sample names: suffix ends with C refers to clinical (CM) and that ends with H refers to healthy (H) milk samples.

### Microbial metabolic functions associated with CM

MR simultaneously analyzed and compared the taxonomic compositions and functional profile in our metagenomic sequences in several ways. On average, the putative genes with known predicted protein and known functions were 3.94% and 5.51%, respectively suggesting that a large proportion of the genes encoding for different functional properties are yet unknown (Supplementary Data 1). By comparing the number of genes assigned to each KEGG pathway between the groups, we found a series of significant differences (*p*=0.001) that lead to the functional divergence among CM and H milk microbiotas. The PCoA analysis at level 3 subsystems showed that CM metagenome separately distributed from H milk metagenome indicating significant functional differences (*p*=0.035) (Supplementary Fig. 4). In the comparative analysis, we found that genes associated with metabolism (central carbohydrate, amino acids, cofactors, vitamins, prosthetic groups and pigment), substrate dependence, clustering-based subsystems, cell motility (bacterial chemotaxis, flagellar assembly, invasion of epithelial cells), phases, prophages, transposable elements and plasmids, regulation and cell signaling, stress response, virulence, disease and defense, immune and infectious diseases, cancer pathways were significantly (*p*< 0.05) over represented and positively correlated with bovine CM (Fig. 6, 7, Supplementary Data 3).

**Fig. 6.**
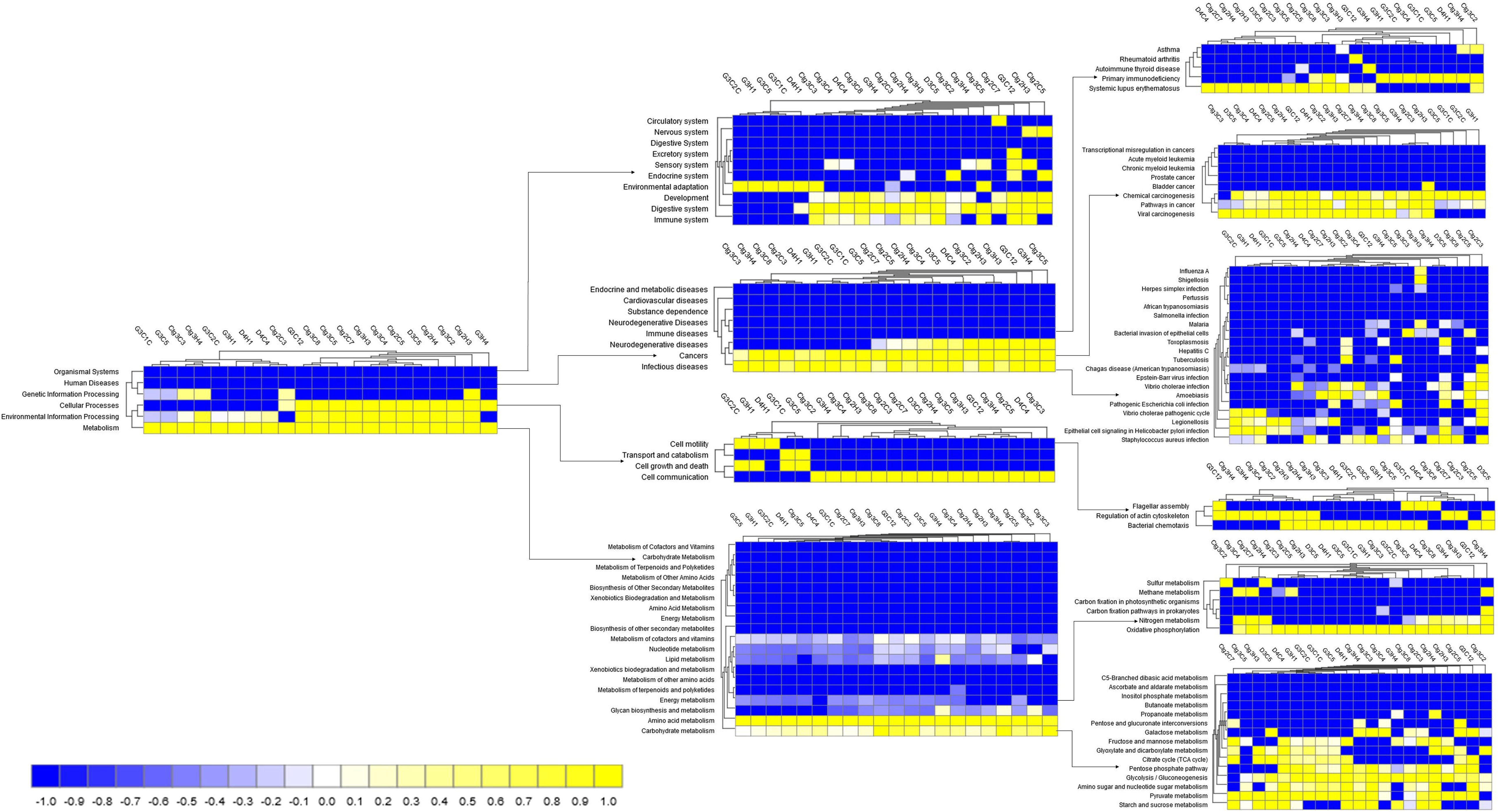
Shotgun whole metagenome sequencing (WMS) reveals differences in functional microbial pathways. Heatmaps show the average relative abundance hierarchical clustering of the predicted KEGG Orthologs (KOs) functional pathways of the microbiota across all samples. The color bar at the bottom represents the relative abundance of putative genes. The color codes indicates the presence and completeness of each KEGG module, expressed as a value between −1 (low abundance), and 1 (high abundance). The greener colors indicates the more abundant patterns, whilst redder cells accounts for less abundant, and the dark black cells represents the absence of the KOs in that particular sample. Sample name: suffix ends with C refers to clinical mastitis (CM) and that ends with H refers to healthy (H) milk samples.

**Fig. 7.**
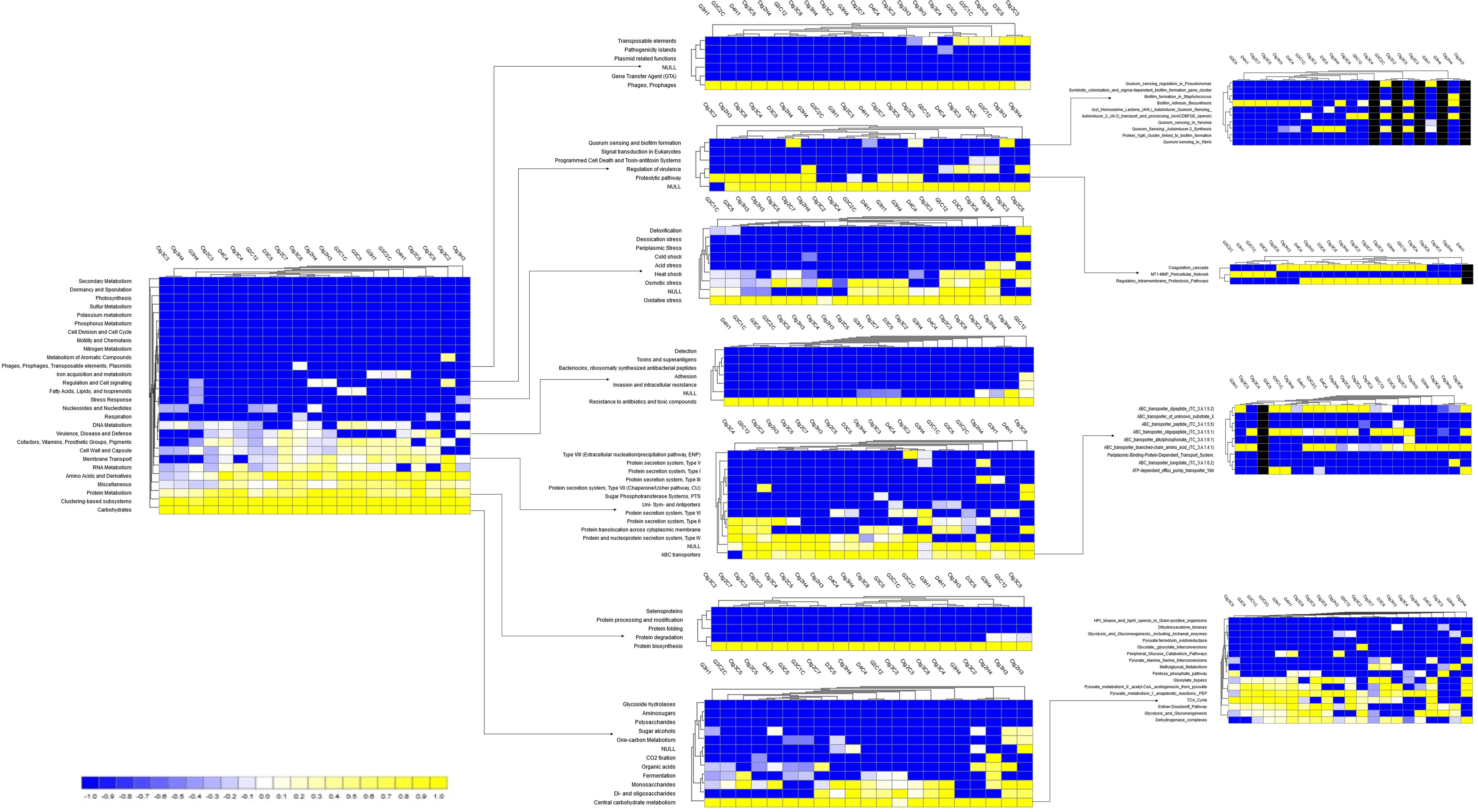
Functional annotation using the SEED subsytem definition. Comparison of metagenomic profiles at the SEED subsystem level 3. Only a selection of subsystems showing significant differences between the two sample groups is shown. The subsystems less abundant in a given metagenome are shown in blue, and more abundant subsystems are represented in yellow colors. The color codes indicated the presence and completeness of each subsystem module, expressed as a value between −1 (low abundance), and 1 (high abundance). The color bar at the bottom represents the higher relative abundance of putative genes. Sample name: suffix ends with C refers to clinical mastitis (CM) and that ends with H refers to healthy (H) milk samples.

Genes associated with citrate synthase (CS, *glt*A), fumarate hydratase class I (*fum*A, *fum*B), oxidative phosphorylation, bacterial translation, ribosome biogenesis and tRNA amino-acylation were significantly enriched in the metabolic pathways of CM associated microbiomes. The CM associated microbiotas had significantly (*p*<0.001) higher relative abundance (50.51%) of genes coding for benzoate degradation than the H milk biomes (36.41%). The CM milk microbes had upregulation of genes for energy metabolism including one carbon metabolism, sulfur and methane metabolism than H milk microorganisms. The relative abundance of genes encoding ABC transporter (38.97%) and bacterial chemotaxis (68.61%) remained significantly higher in CM microbes than those detected in H milk biomes (*p*<0.005). Among the pathways in infectious diseases, genes coding for epithelial cell signaling, epithelial cells invasion, Legionellosis, *Vibrio cholerae* pathogenic cycle, *Staphylococcus aureus*, *Salmonella* and pathogenic *Escherichia coli* infection were mostly abundant in CM metagenome. Likewise, there was a predominant abundance of genes responsible for glutathione S-transferase (GST), breakpoint cluster region protein (BCR1), fumarate hydratase class II (*fum*C), and pyruvate kinase (pk) in different pathways causing mammary gland cancer. We observed significantly higher abundance of genes encoding apoptosis in CM microbiomes, and in contrast, the relative abundance of proteins that are involved in various cellular functions (cell growth and differentiation) through the p53 signaling pathway remained higher in H milk (p<0.05). The CM milk microbiomes had significantly (*p*< 0.001) higher number of reads (64.29%) coding for severely combined immune deficient gene adenosine deaminase (ADA) than H milk microbes (28.58%) (Supplementary Fig. 5). Furthermore, sporulation related hypotheticals and CRISPR-associated proteins (*Cas*1, *Cas*2, and *Cas3)* remained higher in CM metagenomes than H milk microbes (Supplementary Data 3).

We found that the CM microbiotas had significantly higher abundance of genes encoding for oxidative stress (36.46%), pathogenicity islands (10.13%), phage related transposable elements (19.48%), phage packaging machinery (6.37%), phage replication (6.70%) and phage regulatory gene expression (7.10%) than those of H milk biomes (*p*< 0.003). However, the phage lysogenic conversion related genes remained higher in abundance among the healthy milk microbes. A deeper look at microbial genes associated with regulation and cell signaling revealed that CM microbes had significantly higher expression of this gene compared to healthy milk microbiotas (*p*=0.001). Within this subsystem, genes coding for two-component regulatory system BarA-UvrY (*Sir*A; CM= 85.78% vs H= 67.41%), pericellular trafficking and cell invasion- the membrane type-1 matrix metalloproteinase (MT1-MMP; CM= 86.59% vs H= 73.80%), programmed cell death (CM= 55.00% vs H= 28.57%), and intra-membrane regulatory proteolytic pathway- endoplasmic reticulum chaperon *grp*78 (BiP; CM= 92.85% vs H= 71.42%) were predominantly found to be associated with the onset of bovine CM. We also identified novel associations of biofilm formation (BF) properties among the microbes identified in both metagenomes. The relative abundance of genes coding for protein *Yjg*K cluster linked to biofilm formation, biofilm PGA synthesis, deacetylase *Pga*B, N-glycosyltransferase *Pga*C, and auxiliary protein *Pga*D had statistically over expression among mastitis causing pathogens (*p*=0.035). In contrast, the genes coding for quorum sensing (QS) in particular to QS in *Yersinia*, *Pseudomonas* and *Vibrio* remained over expressed in H milk metagenomes. Moreover, of the assigned reads to different levels SEED subsystems (6.45 million), 2.63% mapped against 30 and 28 different resistance to antibiotic and toxic compounds (RATC) genes respectively in CM and H milk metagenomes (Fig. 8, Supplementary Data 3). Among them, genes encoding multidrug resistance to efflux pumps, cluster (*mdt*ABCD), operon (*Cme*ABC) and MAR locus, methicillin resistance in *Staphylococci*, vancomycin resistance, arsenic and chromium compounds resistance had two-fold higher relative abundances in CM microbiotas than H milk biomes. There was 5 to 7-fold over expression of multidrug resistance to MAR locus and mercury resistance genes in CM microbes than H milk organisms. In addition, CM causing microorganisms harbored two additional resistance genes; multidrug resistance to operon (*mdt*RP) and aminoglycoside adenyltransferase (Supplementary Data 3).

**Fig. 8.**
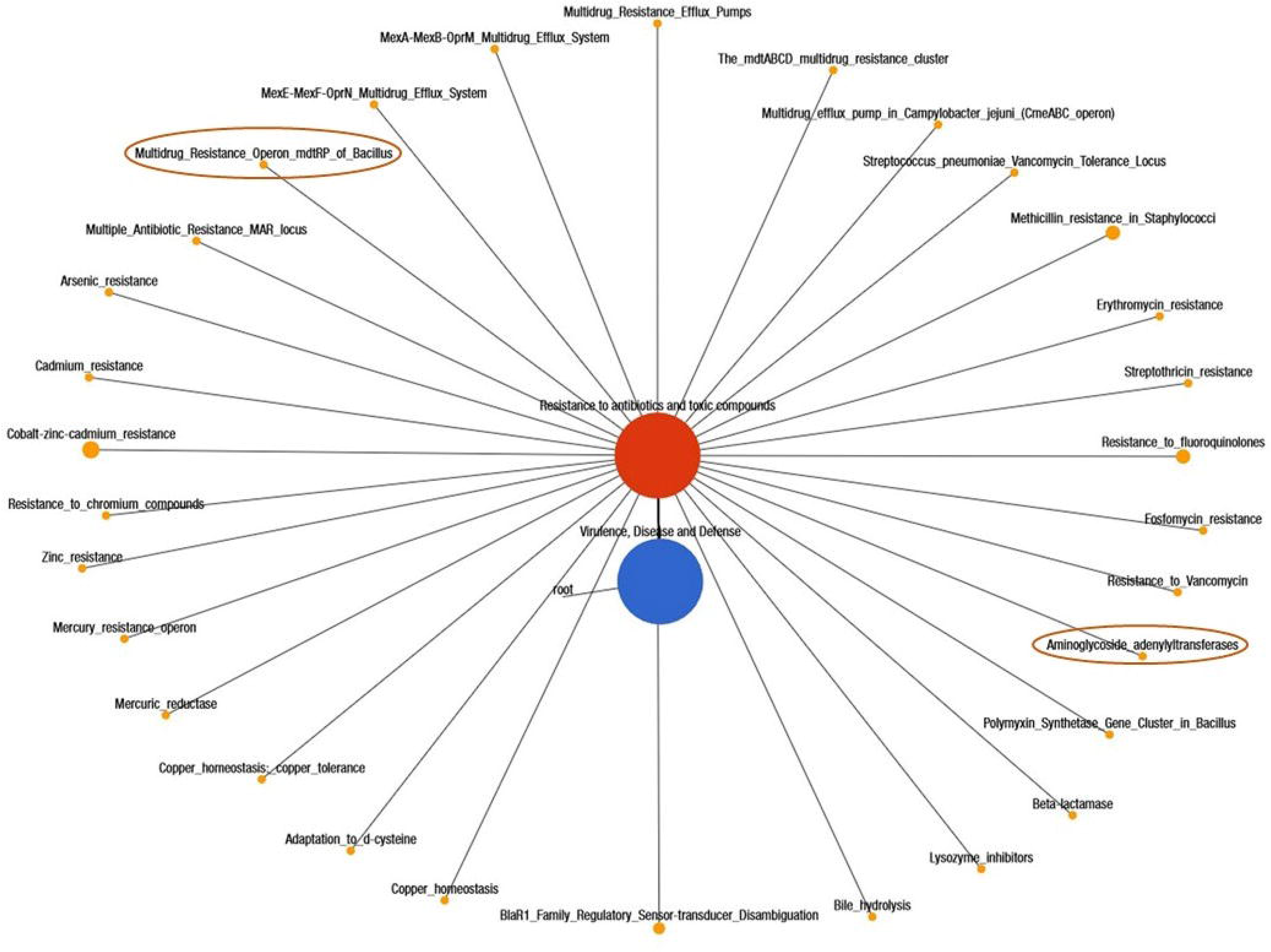
Networks showing distribution of the antibiotics and toxic compounds resistance genes in clinical mastitis and healthy milk samples as detected in subsystem level 3 by using Euclidean distances in MG-RAST. A total of 30 genes in clinical mastitis and 28 in healthy milk metagenomes have been detected. Black lines with yellow circles demarcate the distribution of the resistant genes according to their class across the both metagenomes. The diameter of the circles indicates the relative abundance of the respective genes in both clinical mastitis and healthy milk samples. The two differentially expressed genes (multidrug resistance to operon, *mdt*RP and aminoglycoside adenyltransferase) in clinical mastitis are highlighted in deep yellow circles.

## Discussion

During the last decade metagenomics has helped to shed some light onto the “known unknown” component of the milk microbiome and to enable insights into its taxonomic composition, dynamics, and importance to cows udder health homeostasis. Metagenomic deep sequencing (WMS) of bovine milk has uncovered previously overlooked microbial populations of high complexity with potential roles in regulation of overall microbiome composition and their functional attributes, and in the onset, progression, and treatment strategies of bovine CM. Yet today, 16SrRNA gene sequencing remained as the key approach for studying milk microbiomes, and findings are mostly limited to bacterial identification at the genus level^5, 9, 17^, though this method has serious inherent limitations^18^. However, little is known about the association of other microbes (archaea and viruses), microbiome shift, and particular functional changes during the progression of the disease. The noteworthy findings of the present WMS study are the taxonomic profiling of bacteria at both the species and/or strain-level, the possible association of the archaeal and viral fractions with bacterial mastitis, and the crosstalk between the identified microbiomes and their functional genomics in the association of bovine CM.

The findings generated by PS and MR are much higher in taxonomic resolution and predicted proteins functions, and are consistent with previous 16SrRNA gene based studies^1, 9, 17^. The core bacteria associated with bovine CM such *Acinetobacter, Pseudomonas, Klebsiella, Escherichia, Enterobacter, Staphylococcus, Streptococcus, Bacillus, Pantoea, Shewanella, Ralstonia* etc. remained consistent in both analytic tools although their relative abundances varied even within same sample group. The study demonstrated that in spite of having relatively higher taxonomic abundance, the CM associated microbiomes fluctuate more within the samples than those identified in H milk metagenome corroborating several recent findings^5, 17–19^. To date, around 50 bacterial genera have been reported in bovine milk through 16SrRNA-based metagenomics^1, 9, 17, 20^, while our current WMS study detected 356 and 251 bacterial genera in CM and H milk, respectively indicating the increased discriminatory power of this cutting-edge technology in identifying microbiomes^10, 15^. The observed increase in phylum-level signature of Proteobacteria, Bacteroidetes, Firmicutes and Actinobacteria in CM milk independent of quarter, parity, and breeds of the cows is almost consistent with many of the previous studies^5, 9, 17, 26^. Furthermore, the CM milk metagenome had an inclusion of 68.04% previously unreported bacterial species, most of which are opportunistic in nature. Before now, no substantial information is available regarding the association of different strains of *Acinetobacter* with bovine mastitis, which are opportunistic pathogen to causing CM by acquiring exogenous DNA from other bacteria through horizontal gene transfer, and concomitantly becomes a reservoir of resistant genes^23^. In a recent study, highest association of *Acinetobacter* causing bubaline CM^7^ has been reported supporting our present findings. The H milk metagenome had higher relative abundance of soil or environment (*Micromonospora*) and animal skin (*Pseudomonas*) associated bacteria, which can act as potential opportunist by attacking weak or injured tissues of teats or mammary glands^25, 26^, and can cause acute and/or chronic mastitis^27^. Furthermore, the predominantly identified CM associated bacteria, *Klebsiella pneumoniae* is an opportunistic environmental pathogen, and transmission of this bacterium might occur from contaminated feces and bedding materials^28^ to help in causing CM in healthy mammary glands and/or quarters. Gut microbiome plays a key role in maintenance of nutrition, host defense and immune development^29^, and we revealed a close association between gut microbiota and milk microbes in the pathogenesis of bovine CM^30^. Additional support for this finding includes, the potential existence of an endogenous entero-mammary pathway, through which gut bacteria migrate to the mammary gland, and this could explain the predominating presence of gut bacteria such as phyla Proteobacteria, Bacteroidetes, Firmicutes, Actinobacteria, Fusobacteria, and Tenericutes, with *Acinetobacter*, *Campylobacter*, *Bacillus*, *Enterobacter*, *Staphylococcus*, *Streptococcus, Kocuria* genera in CM milk^30–32^. These pathogens use very efficient strategies to evade host defenses in order to colonize and invade mammary tissues through adhesion^33^, thereby damage host cells and fight with cows immune systems to producing clinical and/or chronic mastitis^33–35^. Despite relatively lower abundance, the rest of the bacterial genera, species/strains detected across the clinical milk metagenome had symbiotic network, and positive correlation with CM. Our study marks an additional step towards identifying the significant co-occurrence of archaea and viruses with bacterial population in bovine milk. Unlike bacteria, the relative abundance and diversity of archaea^36^ and viruses^37^ remained substantially lower. Currently there is no extensive evidence supporting the role of archaea and viruses in the pathogenesis of bovine mastitis, however these microbes mostly cease the opportunity during the pathophysiological changes in the mammary glands created by bacteria^38^. The full spectrum of outcomes from these archaea-cows interactions, whether it altered host mammary gland physiology remained as a mystery. Thus, it is hypothesized that archaea might follow the exact mechanisms of bacterial pathogens producing bovine CM^36^. Most of the detected viral genera belonged to the order Caudovirales which consists of the three families of tailed bacterial viruses (bacteriophages) infecting bacteria and archaea. The host range of Caudovirales is very broad and includes all major bacterial phyla found in both metagenomes: Firmicutes, Bacteroidetes, Proteobacteria, and Actinobacteria. This corresponded with an increase relative abundance of these bacterial taxa in CM milk samples together with an over presentation of Caudovirales taxa compared with H milk metagenome^39^. In addition, we revealed the association of Herpesvirales (*Macavirus* and *Rhadinovirus* genera) with bovine CM^39, 40^. Our current findings demonstrated that viruses neither cause bovine mastitis directly nor play role in the initiation of the disease process, but later, when bacterial infection of the udder occurs, they replicate in the immune and epithelial cells of the udder and/or milk ducts, and may act as predisposing factor as well as primary etiological agent for more severe and prolonged mastitis^41^.

The KEGG pathways and SEED subsystems of MR pipeline uncovered significant differences in microbial metabolic functions in both metagenomes^5, 42^ as supported by several previous reports on mastitis in lactating cows^9^ and women^5^. The CM microbiota had significantly higher abundance of Proteobacteria and Bacteroidetes, which are well-known bacteria to utilizing milk oligosaccharides; one carbon metabolism^43^. Genes associated TCA cycle (*glt*A, *fum*A) and energy metabolism (oxidative phosphorylation) remained over expressed in CM microbiomes, which might be associated with host-pathogen interactions during the progression of bovine mastitis^44, 45^. Increased benzoate degradation by different strains of *Acinetobacter* and *Klebsiella* in CM metagenome through TCA cycle is thought to promote bacterial growth and virulence factors expression during pathogenesis^46, 47^. To elucidate the role of bacterial chemotaxis in bovine mastitis, we found that genes coding for bacterial chemotaxis is predominantly abundant in CM milk microbiomes suggesting their role in early phase of mastitis for attachment to or entry into the udder tissues and virulence regulation^48^. The p38 signaling pathway exerts its biological effects in the pathophysiology of bovine CM through several complex biologic processes including expression of many cytokines, transcription factors, cell surface receptors, enzymes and oxidative stress mediators^49^. The p38 mediated transcriptional regulation of matrix metalloproteinase-2 (MMP-2)^50^ and pro-inflammatory mediator cyclooxygenase-2 (COX2)^49^ can potentially contribute to mammary gland cancer and/or glandular mastitis. The up-regulation of genes coding for programmed-cell death during host–pathogen interactions in CM is associated with increased secretion of bacterial toxins, or pro-inflammatory mediators^51^. Diverse groups of microbiomes (bacteria and viruses) causing bovine CM might induce cell death with their apoptotic features^51^. The predominantly identified membrane type-1 matrix metalloproteinase (MT1-MMP) across the CM metagenome is a pro-invasive protease regulating various cellular functions, macrophage migration to the inflamed mammary tissues, and causes adenocarcinoma in cows udder^52^. We demonstrated that endoplasmic reticulum chaperon (GRP78/BiP) associated gene signatures are highly expressed in CM microbiotas, which can promote tumor proliferation and metastasis in mammary tissues^53^. Biofilm formation can be a strain specific or genetically linked trait, representing a selective advantage in pathogenesis of mastitis. The relatively over expression of genes encoding protein YjgK cluster linked to biofilm formation, and biofilm PGA synthesis in CM microbiomes is in accordance with several earlier reports^54^. Moreover, biofilm formation can also be harmful to host tissues since they can promote the phagocyte release of lysosomal enzymes, proliferation of reactive oxygen and nitrogen species, and transfer of antibiotic resistance^55^. The observed increase abundance of genes for primary immune diseases; adenosine deaminase (ADA) in CM pathogens is responsible for inhibition of T cell maturation and lymphocytic proliferation^56^, very low CD4 count^57^, cell-to-cell communication^58^, and therefore could be used as a selective marker for bovine CM diagnosis. CRISPR/Cas systems are present in both pathogenic and commensal organisms found in bovine milk, and play critical roles during the pathogenesis of mastitis by evading the hosts defense system particularly under stress condition^59^. The type III and IV secretion systems found on the pathogenicity islands of CM associated microbes are capable of producing immunosuppression in cows by delivering effector proteins^960^. Phages, which are the regulators of bacterial population, play important and diverse roles in all bacterial ecosystems^61^, but their precise impact on the milk microbiota is far from being understood. The relatively over presentation of genes coding for phage related transposable elements, phage packaging machinery, phage replication and phage regulatory gene expression in CM microbes may suggests that bacteriophages participate in the horizontal gene transfer among the members of bovine milk microbiomes, and ultimately to mammary gland pathogens^39^. We propose that as obligate parasites bacteriophages naturally found in raw milk, replicate in bacterial host, follow the lysogenic cycle, disrupt host metabolism and, ultimately, causing death of bacterial cell during the immunosuppression states of the cows, and finally release new phage particles^39, 40^.

Bovine milk microbiomes are a wide source of resistance to antibiotic and toxic compounds (RATC) genes and the pathogenic bacteria within this potential reservoir are becoming more resistant. The current metagenomic deep sequencing provides a wealth of information not only on RATC genes, but on the entire gene content thereby enabling the identification of the community composition and metabolic profile. We found that all of the samples in both metagenomes harbored RATC genes (2.63%) indicating their wide and indiscriminate use in Bangladeshi dairy farms. However, most of the resistant genes in RATC functional groups remained predominantly higher in CM milk microbes. Although the knowledge on uncontrolled spread of antibiotics resistant genes in bovine mastitis pathogens^62^ are increasing, but information on heavy metal resistance is yet unavailable. This worrisome trend in increasing RATC against mastitis pathogens has become a major concern for the dairy holders of Bangladesh, given the seriousness of such problems; effective therapies using alternative medicines are needed for successful prevention and control of bovine mastitis. The novel WMS technology in combination with improved bioinformatic analysis of milk microbiome identifies the comparative microbial communities associated with bovine CM and H quarters. The significant differences in the microbiome compositions and protein functional diversities in two groups implicated the association in the progression of the pathophysiological conditions of the disease.

## Methods

### Study population and sampling

Details of study population and collected samples are presented in Supplementary Table 1. A total of 21 milk samples (14, CM and 7, H) from 21 lactating crossbred cows at their early stage of lactation (within 10-40 days of calving) were collected from three districts of Bangladesh (Chattagram= 12, Dhaka= 3, Gazipur=6). The sampling patterns followed collection of two CM and one H milk samples from the same farm. Approximately 15-20 ml of milk from each cow was collected in a sterile falcon tube during the morning milking (8.0-10.0 am) with emphasis on pre-sampling disinfection of teat-ends and hygiene during sampling^1, 63^. The milk samples were then transported to the laboratory, and stored at −20°C until DNA extraction.

### DNA extraction and sequencing

Genomic DNA (gDNA) was extracted by an automated DNA extraction platform (Promega, UK) following previously described protocols^5, 16^. DNA quantity and purity was determined with NanoDrop (ThermoFisher, USA) by measuring 260/280 absorbance ratios. Sequencing libraries were prepared with Nextera XT DNA Library Preparation Kit^64^ according to the manufacturer’s instructions, and paired-end (2×150 bp) sequencing was performed on a NextSeq 500 machine (Illumina Inc., USA) at the Genomics Core facility at The George Washington University. Our metagenomic DNA yielded 483.38 million reads with an average of 23.01 million (maximum=35.10 million, minimum=6.77 million) reads per sample (Supplementary Data 1).

### Sequence reads preprocessing

The resulting FASTQ files were concatenated and filtered through BBDuk^13^ (with options k=21, mink=6, ktrim=r, ftm=5, qtrim=rl, trimq=20, minlen=30, overwrite=true) to remove Illumina adapters, known Illumina artifacts and phiX. Any sequence below these thresholds or reads containing more than one ‘N’ were discarded. On an average, 20.16 million reads per sample (maximum=32.33 million, minimum=4.71 million) passed quality control step (Supplementary Data 1).

### Microbiome community analysis

We analyzed the WMS data using mapping-based and assembly-based hybrid methods PathoScope 2.0 (PS)^65^ and MG-RAST 4.0 (MR)^8^. In PS analysis, a ‘target’ genome library was constructed containing all bacterial and archaeal sequences from the NCBI Database (https://en.wikipedia.org/wiki/National_Center_for_Biotechnology_Information) using the PathoLib module. The reads were then aligned against the target libraries using the very sensitive Bowtie2 algorithm^15–16^ and filtered to remove the reads aligned with the cattle genome (bosTau8) and human genome (hg38) as implemented in PathoMap (−very-sensitive-local -k 100 --score-min L,20,1.0). Finally, the PathoID^66^ module was applied to obtain accurate read counts for downstream analysis. In these samples, an average of 12.90 million aligned reads per sample mapped to the target reference genome libraries (96.24 %) after filtering the cow and human genome (Supplementary Data 1). The raw sequences were simultaneously uploaded in MR server (release 4.0) with proper embedded metadata and were subjected to the quality filter containing dereplication and removal of host DNA by screening^67^ for taxonomical and functional assignment.

### Diversity analysis

Alpha diversity (diversity within samples) was estimated using the Shannon index for both PS and MR reads output. To test beta diversity (differences in the organismal structure) of the milk microbiome, a principal coordinate analysis (PCoA) was performed based on weighted-UniFrac distances (for PS data) through Phyloseq R^68^, and Bray-Curtis dissimilarity matrix (for MR data)^69^. In addition, non-metric multidimensional scaling (NMDS) on PS data was also used for beta diversity analysis between the sample groups^70^. Taxonomic abundance was determined by applying the “Best Hit Classification” option using the NCBI database as a reference with the following settings: maximum e-value of 1×10^−30^; minimum identity of 95% for bacteria, 60% for archaea and viruses, and a minimum alignment length of 20 as the set parameters. The phylogenetic origin of the metagenomic sequences was projected against the NCBI taxonomic tree and determined by the lowest common ancestor (LCA) with the same cutoff mentioned above. Two phylogenetic trees consisting of 363 and 146 bacterial strains respectively in CM and H metagenomes with >80% taxonomic identity were constructed using the neighbor-joining method in Clustal W (version 2.1)^71^, and FigTree (version 1.5.1)^13^.

### Statistical analysis

The characteristics of cows with and without CM were compared using Fisher’s exact test for categorical variables, and Mann-Whitney U test for quantitative variables^21^. The Shapiro-Wilk test was used to check normality of the data, and the non-parametric test Kruskal-Wallis rank sum test was used to evaluate differences in the relative percent abundance of taxa in CM and H groups. For the functional abundance profiling, the statistical tests were applied at different KEGG and SEED subsystems levels in MR pipeline. Differences between the pipelines were evaluated using ANOVA and the Friedman rank sum test. A significance level of alpha=0.05 was used for all tests^8^.

## Funding and Acknowledgements

The Bangladesh Bureau of Educational Information and Statistics (BANBEIS), Ministry of Education, Government of the People’s Republic of Bangladesh (Grant No. LS2017313) supported this work. The author M. Nazmul Hoque receives Fellowships from the Bangabandhu Fellowship Trust, Ministry of Science and Technology, Government of the People’s Republic of Bangladesh. The authors also thank Keylie Gibson and Stephanie Warnken, PhD students at the Computational Biology Institute, Milken Institute School of Public Health, the George Washington University, USA for their for technical support in learning basic bioinformatics operations.

## Data availability

The raw sequence files have been submitted to NCBI database under BioProject PRJNA529353, and can be accessed to the reviewers when they ask for it. All other relevant data supporting the findings of the study are available in this article as Supplementary information files, or from the corresponding author on request.

## Author contributions

M.N.H., M. S., A.M.A.M.Z.S. and M.A.H. conceived and designed the overall study, and M.N.H. and R.A.C. carried out laboratory works including DNA extractions, quality control and preparation for sequencing. M.A.H., R.A.C. and K.A.C. contributed reagents/materials/analysis tools and sequencing. M.N.H. and A.I. conceived, designed and executed the bioinformatics analysis and M.N.H. interpreted the results and prepared the manuscript. M.S., K.A.C., M.A.H contributed intellectually to the interpretation and presentation of the results. Finally, all authors have approved the manuscript for submission.

## Competing interests

The authors of this study declare no competing interests.

