## Supplementary figures and images for "Association of milk microbiome in bovine clinical mastitis and their functional implications in cows in Bangladesh"

### Supplementary Fig. 1

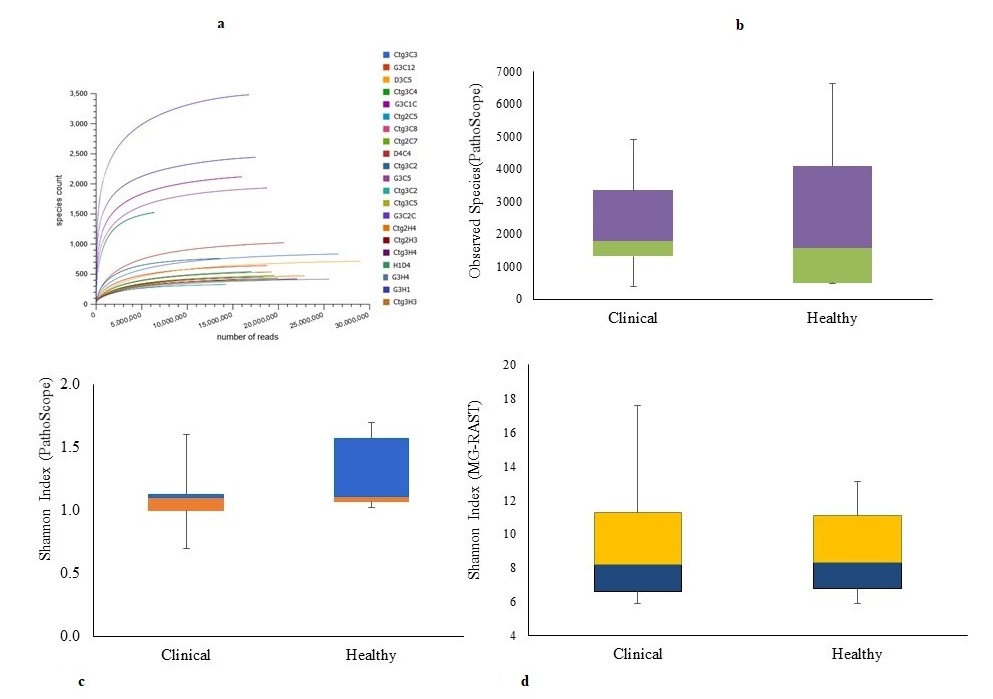

### Supplementary Fig. 2

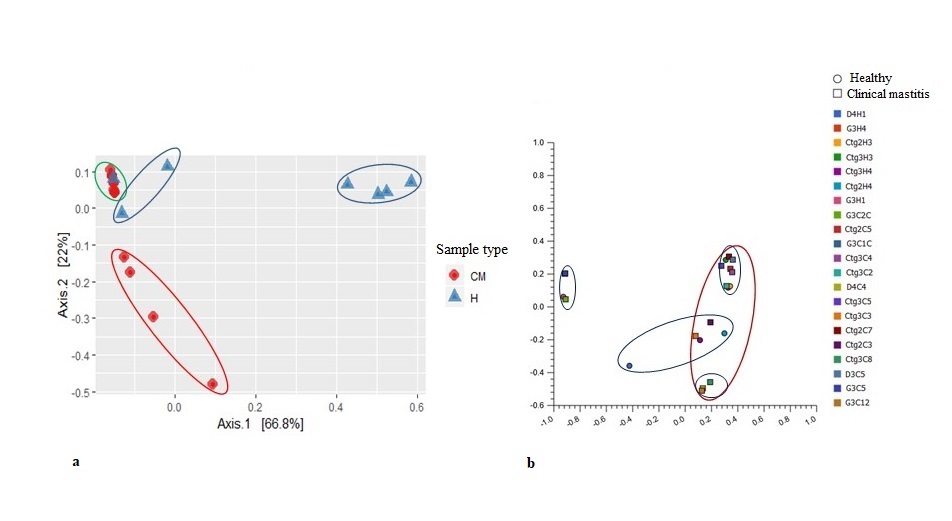

### Supplementary Fig. 3

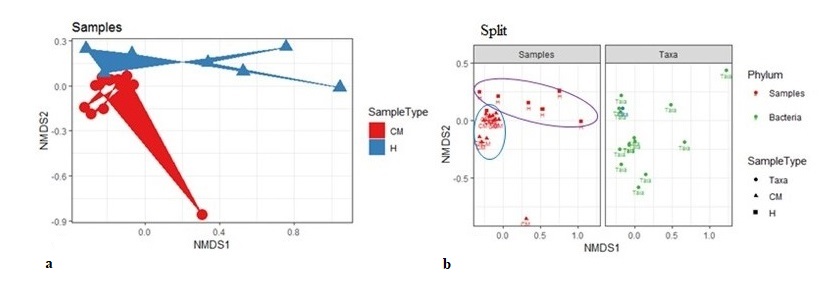

### Supplementary Fig. 4

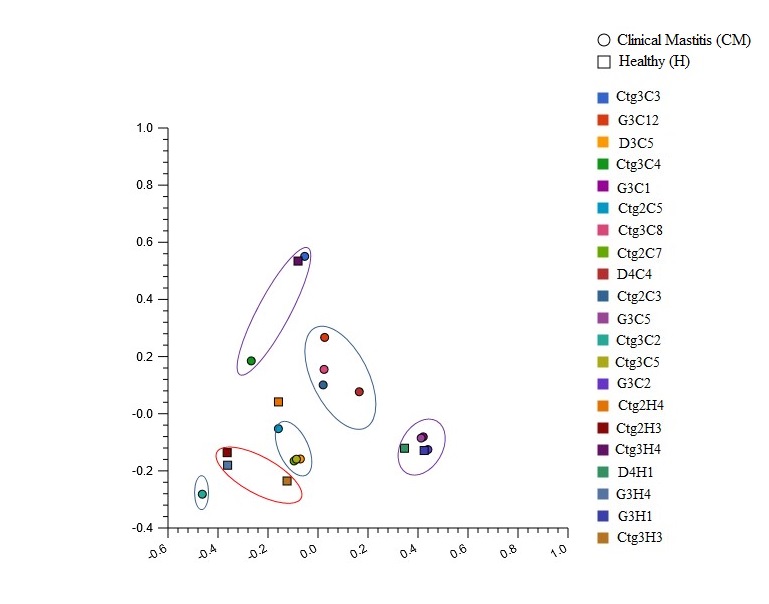

### Supplementary Fig. 5

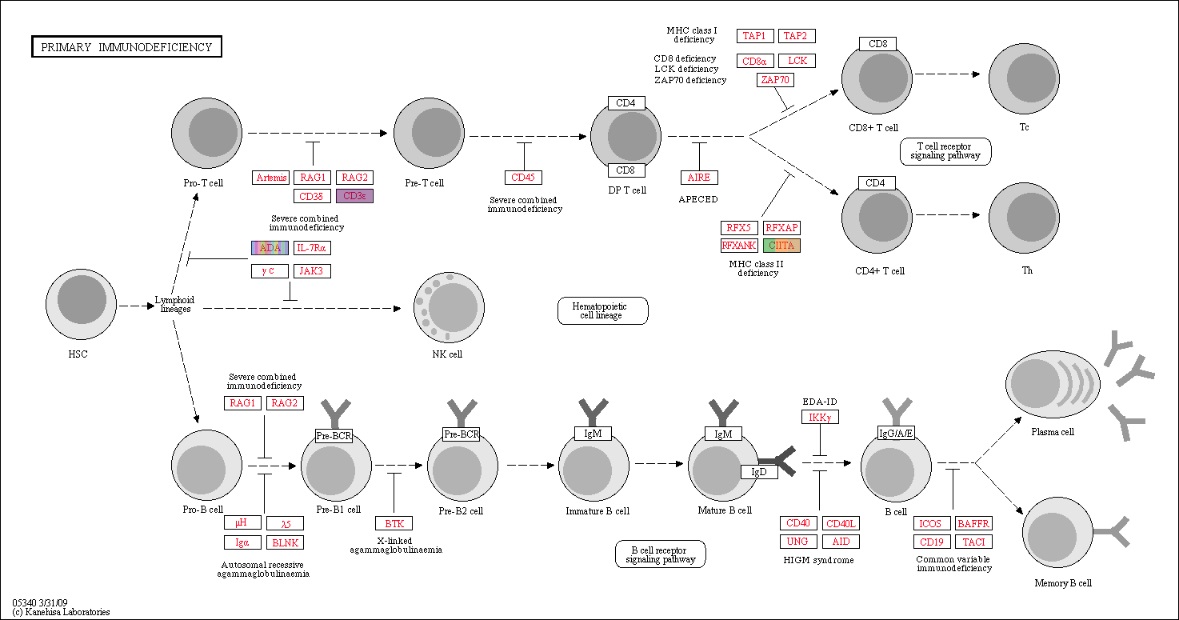
