## Supplementary Fig. Legends for "Association of milk microbiome in bovine clinical mastitis and their functional implications in cows in Bangladesh"

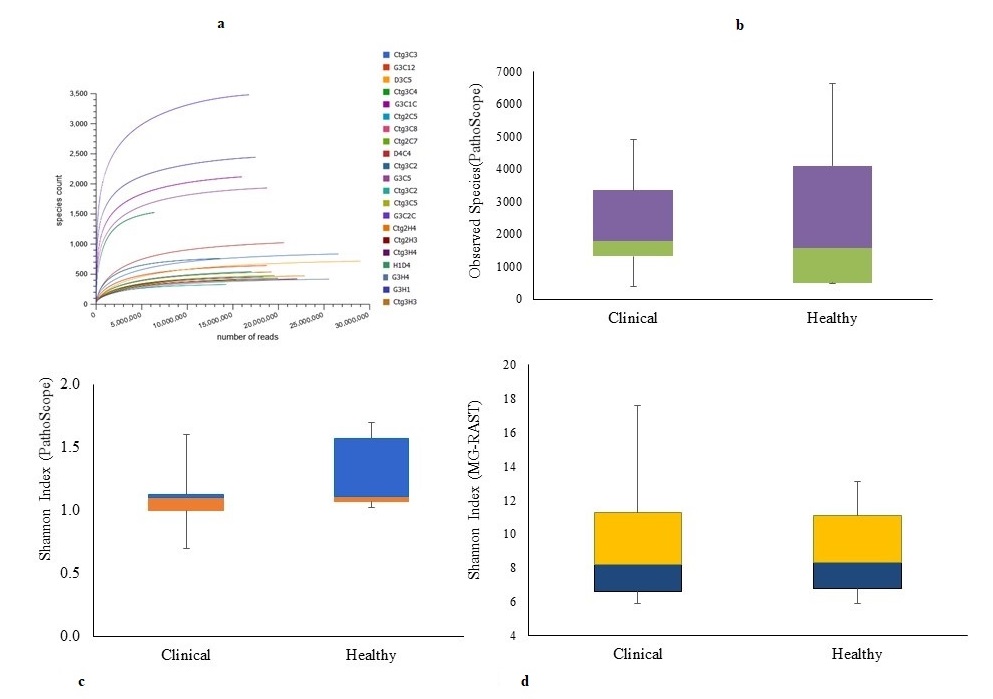

**Supplementary Fig. 1** Alpha-diversity analysis through observed species and Shannon diversity indices in clinical mastitis (CM) and healthy (H) milk samples. **a** Top Left: Rarefaction curves indicate the number of detected species (y axis) per sample (x axis), and **b** Top Right: Alpha-diversity using observed species richness, **c** Bottom Left: & **d** Bottom Right: Alpha-diversity calculation with Shannon diversity index. The observed species richness (P_Observed_ = 0.013) and Shannon diversity (P_Shannon_ = 0.001) in both PS and MR remain significantly higher for H milk metagenomes compared to CM milk metagenomes. The rarefaction curve represented the richness of the observed species and indicated that the sequencing depth was sufficient to fully capture the diversity as existed. Compared to H metagenome, microbiome of CM had significantly reduced Shannon-estimated microbial richness (Mann Whitney U test, p=0.005 and 0.007, respectively). The boxplots showing the species level Shannon diversity of the WMS reads at 25^th^, 50^th^, and 75^th^ percentiles had a shift towards decreasing richness in relation to cows mastitis condition (P_Shannon_ = 0.001), but not for the healthy states of the udder. Minimum and maximum values for each species, as computed by PS and MR are shown as whiskers. In addition, species richness in CM and H milk metagenome also significantly differed between the tools used for diversity analysis (Mann Whitney U test, p=0.039 and 0.001 for PS and MR, respectively). Sample names: suffix ends with C refers to clinical (CM) and that ends with H refers to healthy (H) milk samples.

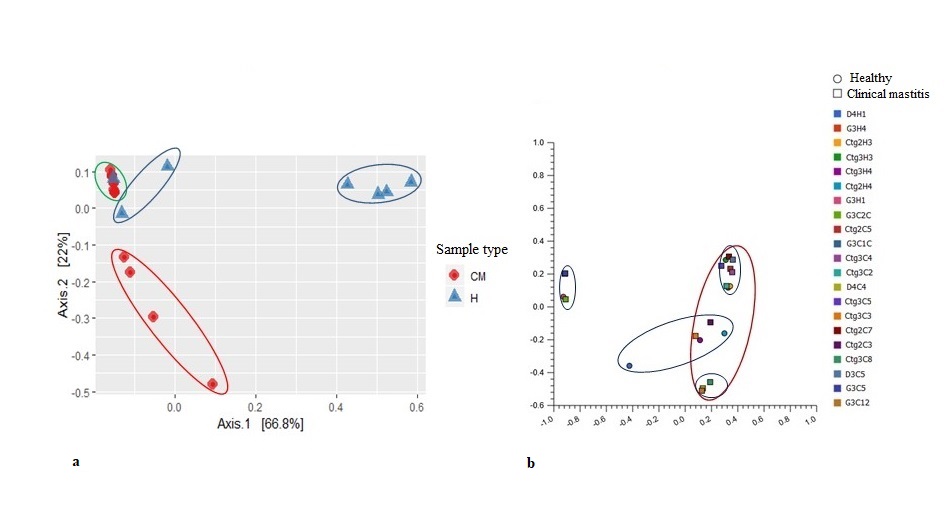

**Supplementary Fig. 2** The beta diversity (Principal coordinate analysis; PCoA) of microbial communities in CM and H milk sample pools measured on- **a** Left: weighted-UniFrac distance metrics (strain level), and **b** Right: Bray-Curtis distance method (genus level). The Kruskal–Wallis and Mann-Whitney tests shows variations in microbial compositions. The PCoA plot on weighted-UniFrac distance metrics at strain level (for PS data) reveals that most of the CM (green circle) and H (blue circles) samples appear more distantly with some sub-clustering (red and blue circles) indicating significant group differences (*p*=0.001). This microbial community structure differences among the samples of the both groups could be explained by a large percentage of variation in the first (66.8%) and second (22.0%) axes. On the other hand, PCoA plot based on the Bray-Curtis distance method at genus level (for MR data) shows that majority of samples in both groups appear more closely (pink circle) with several intermixed sub-clustering (light blue circles) of the microbiotas in both sample groups representing less diversity differences. Sample names: suffix ends with C refers to clinical (CM) and that ends with H refers to healthy (H) milk samples.

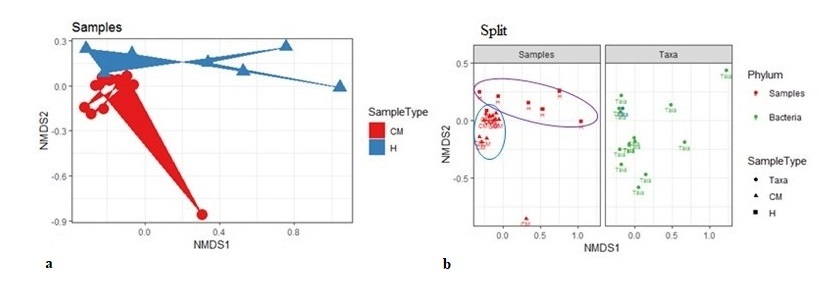

**Supplementary Fig. 3** The NMDS ordination plot shows a clear separation of CM and H milk samples by weighted-UniFrac distance on PS data at phylum level. **a** Left: Clinical mastitis (CM) associated microbial communities appeared to cluster towards the left bottom side of NMDS1 plot while non-clinical healthy milk (H) associated communities cluster towards the opposite side of NMDS1(top). **b** Right: Microbiome associated with CM appear to closely cluster towards the left side of NMDS1 (blue circle), and H milk microbiota diversely clustered towards the opposite side of NMDS1 (pink circle). Thus, the phylum level microbiomes signature on PS data between CM and H milk revealed distinct separation similar to the differences found at strain level between the groups of both sample categories. Statistical analysis using Kruskal–Wallis and Mann-Whitney tests showed significant microbial diversity variation in both group of samples in PS and MR (*p*= 0.001).

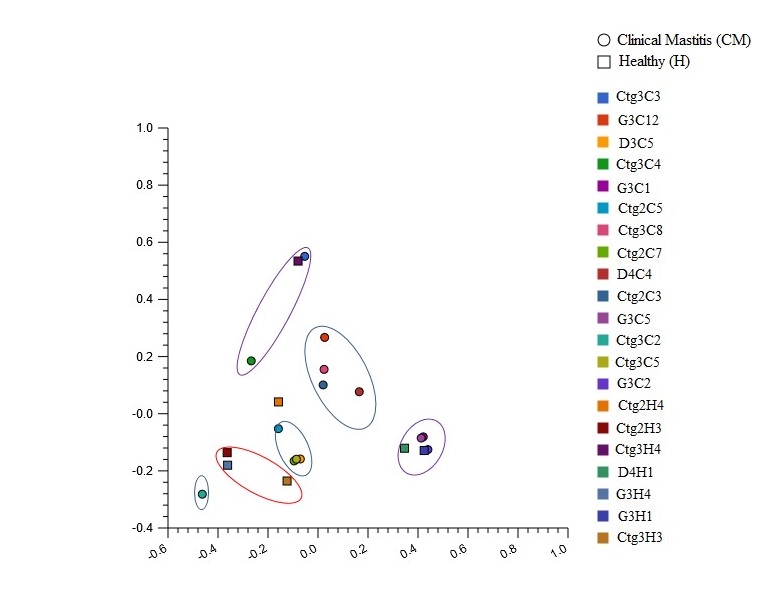

**Supplementary Fig. 4** The beta diversity analysis (PCoA) of functional properties of the microbial communities in clinical mastitis (CM) and healthy (H) milk samples at SEED subsystem level 3. The PCoA plot of milk microbial communities and their functional attributes based on Bray-Curtis distance method showed clear separation between the CM and H milk biomes with several subclustering indicating significant differences in functional attributes of the microbiomes of both groups. Statistical testing of variation in microbial functional attributes was carried out using Kruskal–Wallis (*p*=0.035). Sample names: suffix ends with C refers to clinical (CM) and that ends with H refers to healthy (H) milk samples.

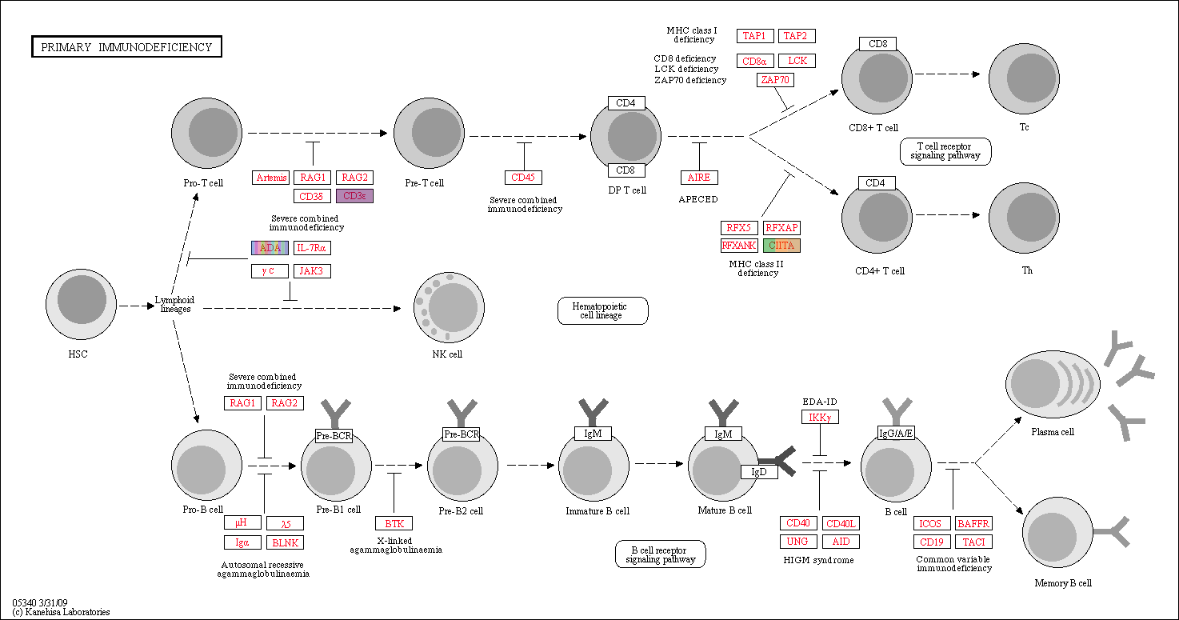

**Supplementary Fig. 5** Projection of the metagenome onto KEGG pathways. Colored lines indicated pathways whose activity was highly expressed in the both metagenomes. The genes associated with adenosine deaminase (ADA) deficiency were highly expressed in clinical mastitis metagenomes while the class II major histocompatibility complex (CIITA) and cluster of differentiation 3 (CD3) gene expression were high in healthy milk microbiomes. The data were analysed with the MG-RAST server using the KEGG mapper for functional analysis with a maximum e-value of 1e^-30^, a minimum identity of 60%, and a minimum alignment length of 20 amino acids in protein databases.
