## Supplementary Table 1 for "Association of milk microbiome in bovine clinical mastitis and their functional implications in cows in Bangladesh"

**Supplementary Table 1:** A total of 21 lactating crossbred cows (including 14 clinical mastitis and 7 healthy) were selected for the study from three different districts of Bangladesh. The cow, farm, milk sample and reads obtained from whole metagenome sequencing (WMS) of each sample related information are given in the table below-

| **Cow ID** | **Sample ID** | **Farm Location** | **Farm ID** | **GIS (Longi-/Latitude)** | **No. of sample/farm** | **Clinical/Healthy** | **Breed** | **Lactation (Days after calving)** | **Parity** | **Reads/sample (Before QC)** | **Reads/sample (after QC)** |
| --- | --- | --- | --- | --- | --- | --- | --- | --- | --- | --- | --- |
| Cow 1 | Ctg2C3 | Chattagram | Ctg1 | 22.20 N 91.98 E | 2 | CM | SW | 15 | 4 | 8885768 | 15601568 |
| Cow 2 | Ctg2C5 | Chattagram | Ctg1 | 22.20 N 91.98 E |  | CM | RCC | 12 | 2 | 20745914 | 19181254 |
| Cow 3 | Ctg2C7 | Chattagram | Ctg2 | 22.20 N 91.98 E | 2 | CM | LZ | 35 | 1 | 23713246 | 21865918 |
| Cow 4 | Ctg3C2 | Chattagram | Ctg2 | 22.20 N 91.98 E |  | CM | XHF | 37 | 3 | 15888132 | 14452552 |
| Cow 5 | Ctg3C3 | Chattagram | Ctg3 | 22.34 N 91.87 E | 2 | CM | XHF | 10 | 2 | 30643824 | 28010688 |
| Cow 6 | Ctg3C4 | Chattagram | Ctg3 | 22.34 N 91.87 E |  | CM | SW | 40 | 2 | 24866832 | 23078692 |
| Cow 7 | Ctg3C5 | Chattagram | Ctg4 | 22.20 N 91.98 E | 2 | CM | RCC | 22 | 5 | 23865184 | 21654684 |
| Cow 8 | Ctg3C8 | Chattagram | Ctg4 | 22.20 N 91.98 E |  | CM | RCC | 18 | 3 | 21767922 | 19655690 |
| Cow 9 | D3C5 | Dhaka | D1 | 23.81 N 90.41 E | 2 | CM | XHF | 20 | 3 | 35106812 | 32334582 |
| Cow 10 | D4C4 | Dhaka | D1 | 23.81 N 90.41 E |  | CM | SW | 35 | 2 | 25456756 | 4710969 |
| Cow 11 | G3C1C | Gazipur | G1 | 24.19 N 90.47 E | 2 | CM | XHF | 24 | 2 | 21526936 | 9636968 |
| Cow 12 | G3C2C | Gazipur | G1 | 24.19 N 90.47 E |  | CM | SW | 18 | 4 | 20090020 | 18727080 |
| Cow 13 | G3C5 | Gazipur | G2 | 24.09 N 90.42 E | 2 | CM | XHF | 13 | 2 | 22980832 | 21332492 |
| Cow 14 | G3C12 | Gazipur | G2 | 24.09 N 90.42 E |  | CM | SW | 32 | 3 | 28792776 | 27018916 |
| Cow 15 | Ctg2H3 | Chattagram | Ctg1 | 22.20 N 91.98 E | 1 | H | RCC | 40 | 3 | 27316282 | 25168186 |
| Cow 16 | Ctg2H4 | Chattagram | Ctg2 | 22.20 N 91.98 E | 1 | H | XHF | 28 | 5 | 28491010 | 26161064 |
| Cow 17 | Ctg3H3 | Chattagram | Ctg3 | 22.34 N 91.87 E | 1 | H | RCC | 36 | 5 | 19692182 | 18189142 |
| Cow 18 | Ctg3H4 | Chattagram | Ctg4 | 22.34 N 91.87 E | 1 | H | XHF | 11 | 2 | 23857586 | 22114556 |
| Cow 19 | D4H1 | Dhaka | D1 | 23.81 N 90.41 E | 1 | H | SW | 27 | 3 | 6772790 | 6009462 |
| Cow 20 | G3H1 | Gazipur | G1 | 24.19 N 90.47 E | 1 | H | XHF | 18 | 3 | 22710894 | 21049986 |
| Cow 21 | G3H1 | Gazipur | G2 | 24.19 N 90.47 E | 1 | H | LZ | 34 | 2 | 30212254 | 27416106 |

*CM, Clinical mastitis milk; H, Healthy milk; SW, Shahiwal crossbred; RCC, Red Chattagram Cattle; LZ, Local zebu; XHF, Holstein Friesian crossbred.
