## Supplementary Table 2 for "Association of milk microbiome in bovine clinical mastitis and their functional implications in cows in Bangladesh"

**Supplementary Table 2:** Taxonomic distribution of bacteria in clinical mastitis (CM) and healthy (H) milk samples by PathoScope (PS) and MG-RAST (MR) analysis.

| **Taxonomic ranks** | **PS** | | **MR** | |
| --- | --- | --- | --- | --- |
|  | CM | H | CM | H |
| Phylum | 8 | 4 | 18 | 12 |
| Class | 17 | 9 | 30 | 22 |
| Order | 44 | 33 | 73 | 53 |
| Family | 90 | 41 | 163 | 124 |
| Genus | 116 | 66 | 359 | 253 |
| Species | 363 | 146 | NA | NA |

*NA: Not detected by the pipeline
