## Supplementary Table 3 for "Association of milk microbiome in bovine clinical mastitis and their functional implications in cows in Bangladesh"

**Supplementary Table 3:** Taxonomic distribution of archaea and viruses in clinical mastitis (CM) and healthy (H) milk samples by MG-RAST (MR) analysis.

| **Taxonomic ranks** | **MR** | | | | |
| --- | --- | --- | --- | --- | --- |
|  | **Archaea** | | **Virus** | | |
|  | CM | H | | CM | H |
| Phylum | 4 | 3 | | 1 | 1 |
| Class | 12 | 11 | | 1 | 1 |
| Order | 16 | 15 | | 3 | 3 |
| Family | 26 | 21 | | 15 | 13 |
| Genus | 54 | 42 | | 35 | 25 |
