## Supplementary Table 4 for "Association of milk microbiome in bovine clinical mastitis and their functional implications in cows in Bangladesh"

**Supplementary Table 4:** Relative abundance of common genera (n=33) found in clinical mastitis (CM) and healthy (H) milk samples by both PathoScope (PS) and MG-RAST (MR) pipelines.

| Genus | **PS** | | **MR** | |
| --- | --- | --- | --- | --- |
|  | Clinical | Healthy | Clinical | Healthy |
| *Acinetobacter* | 41.7616939 | 33.0354368 | 78.4963538 | 72.7709871 |
| *Pseudomonas* | 10.7579069 | 20.2088375 | 17.1658695 | 25.4103685 |
| *Eubacterium* | 9.6293899 | 10.7358556 | 0.0000888 | 0.0000757 |
| *Pantoea* | 0.8852424 | 0.0072491 | 0.4386895 | 0.0361155 |
| *Klebsiella* | 0.6298194 | 0.0003755 | 0.6017880 | 0.0033314 |
| *Psychrobacter* | 0.0055605 | 0.3657184 | 0.0281564 | 0.4859309 |
| *Escherichia* | 0.1359198 | 0.0064116 | 0.4181273 | 0.0261970 |
| *Ralstonia* | 0.0386799 | 0.0451408 | 0.1604339 | 0.1850444 |
| *Salmonella* | 0.0053910 | 0.0028303 | 0.3141617 | 0.0097671 |
| *Streptococcus* | 0.0210769 | 0.0033213 | 0.0964158 | 0.2076828 |
| *Bacillus* | 0.1689121 | 0.1027292 | 0.0206510 | 0.0070414 |
| *Lactococcus* | 0.0037388 | 0.0710469 | 0.0101257 | 0.1885272 |
| *Chryseobacterium* | 0.0305881 | 0.0467292 | 0.0059510 | 0.0622367 |
| *Staphylococcus* | 0.0139171 | 0.0402888 | 0.0182750 | 0.0203670 |
| *Shigella* | 0.0033575 | 0.0000578 | 0.0599545 | 0.0025743 |
| *Corynebacterium* | 0.0024890 | 0.0130542 | 0.0044411 | 0.0376297 |
| *Enterococcus* | 0.0038553 | 0.0046209 | 0.0133676 | 0.0198370 |
| *Kocuria* | 0.0131863 | 0.0200433 | 0.0009104 | 0.0001514 |
| *Lactobacillus* | 0.0034846 | 0.0041011 | 0.0134120 | 0.0087071 |
| *Stenotrophomonas* | 0.0073505 | 0.0002599 | 0.0199848 | 0.0016657 |
| *Listeria* | 0.0032516 | 0.0013285 | 0.0095261 | 0.0096914 |
| *Acidovorax* | 0.0008473 | 0.0005487 | 0.0100590 | 0.0120385 |
| *Macrococcus* | 0.0025525 | 0.0003177 | 0.0063729 | 0.0014386 |
| *Leuconostoc* | 0.0010274 | 0.0020217 | 0.0005995 | 0.0040128 |
| *Delftia* | 0.0022242 | 0.0000866 | 0.0046631 | 0.0006057 |
| *Alcanivorax* | 0.0044696 | 0.0004621 | 0.0019319 | 0.0003029 |
| *Mycobacterium* | 0.0023089 | 0.0014440 | 0.0010214 | 0.0002271 |
| *Clostridium* | 0.0001059 | 0.0000578 | 0.0029755 | 0.0002271 |
| *Prevotella* | 0.0000635 | 0.0001155 | 0.0020429 | 0.0005300 |
| *Rothia* | 0.0003919 | 0.0001733 | 0.0012879 | 0.0003786 |
| *Actinomyces* | 0.0000530 | 0.0000578 | 0.0001554 | 0.0002271 |
| *Cardiobacterium* | 0.0000106 | 0.0000578 | 0.0001110 | 0.0003029 |
| *Brachybacterium* | 0.0000530 | 0.0000578 | 0.0000888 | 0.0002271 |
