## Supplementary Table 5 for "Association of milk microbiome in bovine clinical mastitis and their functional implications in cows in Bangladesh"

**Supplementary Table 4:** Using PathoScope, a total of 247 new or opportunistic bacterial species and/or strains found to be associated with bovine clinical mastitis (CM), and of them most abundant 100 bacterial species and/or strains with their respective relative abundances is given below.

| **SL.NO.** | **New/Opportunistic Species in CM** | **Relative Abundance** |
| --- | --- | --- |
| 1 | *Pantoea dispersa EGD-AAK13* | 0.827 |
| 2 | *Klebsiella oxytoca* | 0.393 |
| 3 | *Kluyvera intermedia* | 0.284 |
| 4 | *Shewanella oneidensis MR-1* | 0.136 |
| 5 | *Kluyvera ascorbata ATCC 33433* | 0.083 |
| 6 | *Klebsiella aerogenes KCTC 2190* | 0.082 |
| 7 | *Kluyvera cryocrescens NBRC 102467* | 0.059 |
| 8 | *Acinetobacter pittii PHEA-2* | 0.057 |
| 9 | *Pseudomonas mendocina ymp* | 0.056 |
| 10 | *Acinetobacter gyllenbergii NIPH 230* | 0.044 |
| 11 | *Enterobacter cloacae subsp. cloacae ATCC 13047* | 0.044 |
| 12 | *[Enterobacter] lignolyticus SCF1* | 0.042 |
| 13 | *Serratia marcescens subsp. marcescens Db11* | 0.040 |
| 14 | *Serratia marcescens FGI94* | 0.038 |
| 15 | *Plautia stali symbiont* | 0.038 |
| 16 | *Kosakonia sacchari SP1* | 0.032 |
| 17 | *Pseudomonas alcaligenes OT 69* | 0.030 |
| 18 | *Enterobacter sp. 638* | 0.029 |
| 19 | *Chryseobacterium haifense DSM 19056* | 0.027 |
| 20 | *Erwinia sp. Leaf53* | 0.024 |
| 21 | *Pseudomonas stutzeri* | 0.023 |
| 22 | *Citrobacter freundii CFNIH1* | 0.022 |
| 23 | *Pseudescherichia vulneris NBRC 102420* | 0.018 |
| 24 | *Pantoea rwandensis* | 0.017 |
| 25 | *Acinetobacter rudis CIP 110305* | 0.014 |
| 26 | *Streptococcus suis BM407* | 0.013 |
| 27 | *Erwinia persicina NBRC 102418* | 0.013 |
| 28 | *Acinetobacter soli NIPH 2899* | 0.013 |
| 29 | *Pluralibacter gergoviae* | 0.013 |
| 30 | *Pseudomonas knackmussii B13* | 0.013 |
| 31 | *Acinetobacter beijerinckii CIP 110307* | 0.011 |
| 32 | *Yokenella regensburgei ATCC 43003* | 0.010 |
| 33 | *Atlantibacter hermannii NBRC 105704* | 0.010 |
| 34 | *Siccibacter turicensis LMG 23730* | 0.009 |
| 35 | *Aeromonas hydrophila SSU* | 0.008 |
| 36 | *Aeromonas hydrophila subsp. hydrophila ATCC 7966* | 0.008 |
| 37 | *Pseudomonas azotifigens DSM 17556* | 0.008 |
| 38 | *Vibrio cholerae O1 biovar El Tor str. N16961* | 0.007 |
| 39 | *Acinetobacter sp. NIPH 298* | 0.007 |
| 40 | *Cronobacter sakazakii* | 0.007 |
| 41 | *Xanthomonas sp. Mitacek01* | 0.007 |
| 42 | *Aeromonas hydrophila YL17* | 0.006 |
| 43 | *Pantoea vagans* | 0.006 |
| 44 | *Cedecea neteri* | 0.006 |
| 45 | *Erwinia typographi* | 0.006 |
| 46 | *Dickeya dadantii 3937* | 0.005 |
| 47 | *Moraxella osloensis* | 0.005 |
| 48 | *Comamonas aquatica* | 0.005 |
| 49 | *Aeromonas veronii* | 0.004 |
| 50 | *Pseudomonas thermotolerans DSM 14292* | 0.004 |
| 51 | *Alteromonas stellipolaris* | 0.004 |
| 52 | *Methylophaga frappieri* | 0.003 |
| 53 | *Vibrio metschnikovii CIP 69.14* | 0.003 |
| 54 | *Escherichia coli IAI39* | 0.003 |
| 55 | *Serratia odorifera DSM 4582* | 0.003 |
| 56 | *Intestinimonas massiliensis* | 0.003 |
| 57 | *Erwinia oleae* | 0.003 |
| 58 | *Spongiibacter tropicus DSM 19543* | 0.002 |
| 59 | *Streptococcus agalactiae 2603V/R* | 0.002 |
| 60 | *Fusobacterium necrophorum HUN048* | 0.001 |
| 61 | *Stenotrophomonas acidaminiphila* | 0.001 |
| 62 | *Agrobacterium tumefaciens* | 0.001 |
| 63 | *Achromobacter xylosoxidans* | 0.001 |
| 64 | *Aeromonas salmonicida subsp. salmonicida A449* | 0.001 |
| 65 | *Porphyromonas levii DSM 23370* | 0.001 |
| 66 | *Arthrobacter sp. Soil736* | 0.001 |
| 67 | *Pantoea sp. PSNIH2* | 0.001 |
| 68 | *Streptococcus plurextorum DSM 22810* | 0.001 |
| 69 | *Pseudomonas fuscovaginae UPB0736* | 0.001 |
| 70 | *Streptococcus entericus DSM 14446* | 0.001 |
| 71 | *Chryseobacterium greenlandense* | 0.001 |
| 72 | *gamma proteobacterium L18* | 0.001 |
| 73 | *Hydrogenophaga intermedia* | 0.001 |
| 74 | *Comamonas testosteroni CNB-2* | 0.0009 |
| 75 | *Shewanella algae* | 0.0009 |
| 76 | *Bacteroides fluxus YIT 12057* | 0.0009 |
| 77 | *Rheinheimera sp. KL1* | 0.0009 |
| 78 | *Erwinia iniecta* | 0.0008 |
| 79 | *Aeromonas simiae* | 0.0008 |
| 80 | *Ochrobactrum pseudogrignonense* | 0.0008 |
| 81 | *Comamonas sp. B-9* | 0.0006 |
| 82 | *Pseudomonas sp. StFLB209* | 0.0006 |
| 83 | *Bradyrhizobium manausense* | 0.0006 |
| 84 | *Bosea thiooxidans* | 0.0005 |
| 85 | *Comamonas kerstersii* | 0.0005 |
| 86 | *Shinella sp. HZN7* | 0.0005 |
| 87 | *Paraglaciecola polaris LMG 21857* | 0.0004 |
| 88 | *Gallaecimonas xiamenensis 3-C-1* | 0.0004 |
| 89 | *Pseudochrobactrum sp. AO18b* | 0.0004 |
| 90 | *Enterococcus columbae DSM 7374 = ATCC 51263* | 0.0004 |
| 91 | *Proteus mirabilis HI4320* | 0.0004 |
| 92 | *Pseudomonas syringae pv. syringae B728a* | 0.0004 |
| 93 | *Elizabethkingia anophelis NUHP1* | 0.0004 |
| 94 | *Franconibacter pulveris DSM 19144* | 0.0004 |
| 95 | *Aeromonas molluscorum 848* | 0.0004 |
| 96 | *Streptococcus dysgalactiae subsp. equisimilis AC-2713* | 0.0004 |
| 97 | *Streptococcus salivarius* | 0.0004 |
| 98 | *Weeksella massiliensis* | 0.0003 |
| 99 | *Weissella confusa* | 0.0003 |
| 100 | *Morganella morganii subsp. morganii KT* | 0.0003 |
